# NetMHCIIphosPan: a machine learning tool for predicting HLA class II antigen presentation of phosphorylated peptides

**DOI:** 10.64898/2026.01.05.697746

**Authors:** Heli M. Garcia Alvarez, Saghar Kaabinejadian, Hooman Yari, Chloe M. Shepherd, William H. Hildebrand, Alessandro Sette, Bjoern Peters, Robert Parker, Nicola Ternette, Morten Nielsen

## Abstract

Phosphorylated peptides presented by human leukocyte antigen (HLA) class II molecules play pivotal roles in immune regulation, yet their characterization and prediction remain challenging due to data noise and limited HLA coverage. Here, we introduce NetMHCIIphosPan, a prediction method for HLA-II antigen presentation of phosphorylated peptides, developed using mass spectrometry (MS)-based immunopeptidomics datasets. Employing a refined peptide identification workflow, we reanalyzed earlier HLA-II phospholigand datasets and trained predictive models, achieving superior performance compared to models trained on original data. Binding motif analysis revealed that HLA-specific preferences for phospholigands closely aligned with those of unmodified ligands. Incorporating unmodified ligands into training further enhanced predictive accuracy, particularly for HLA-DP and HLA-DQ molecules. NetMHCIIphosPan outperformed existing tools, such as NetMHCIIpan-4.3 and MixMHC2pred-1.3, for prediction of HLA antigen presentation of phosphorylated peptides demonstrating robustness and utility. This work establishes NetMHCIIphosPan as a state-of-the-art tool for understanding the HLA-II phospholigandome, with potential applications in immunotherapy and vaccine design.

## Introduction

Major histocompatibility complex (MHC) class II molecules, also known as human leukocyte antigen (HLA) class II in humans, play a central role in the adaptive immune response by presenting peptides to CD4+ T cells. These are heterodimers consisting of α and β chains, with high polymorphism in the β chain only for HLA-DR and in both chains for HLA-DQ and HLA-DP. HLA class II ligands can originate from both exogenous and intracellular proteins processed by the endocytic pathway^1^. This, combined with the locus diversity, allows HLA class II molecules to present a broad range of peptides which is crucial for initiating immune responses and maintaining immune tolerance.

Historically, research on HLA class II antigen presentation has been focused primarily on HLA-DR molecules, given their higher expression^2^ and established links to autoimmune diseases^3^. However, recent advancements in immunopeptidomics and machine learning (ML) have led to an increased understanding of the peptide repertoires presented by all HLA class II loci. These advances include a transformed view of the contribution of HLA-DRB3/4/5 molecules to the overall HLA-DR immunopeptidome^4^, improved predictive accuracy and allele coverage for HLA-DQ and DP loci^5,6^ and confirmation of α and β chain-pairing rules that limit the diversity of HLA-DQ heterodimers^5^. Furthermore, motif deconvolution methods have identified an inverted peptide binding mode, primarily restricted to HLA-DP heterodimers formed by specific HLA-DPA1 molecules^6^. Building on these findings, pan-specific prediction tools such as NetMHCIIpan-4.3 have been developed^6,7^ offering a close to complete understanding of the rules defining HLA class II antigen presentation of unmodified peptides, significantly improving the ability to deconvolute complex immunopeptidomes.

However, post-translational modifications (PTM) play a key role in defining the function of proteins because they regulate activity, localization, and protein-protein interactions. Phosphorylation, a ubiquitous PTM, is known to influence protein structure and regulate protein function^8^. Yet its role in contributing to the HLA-II immunopeptidome is not fully understood. Preliminary earlier studies in mice have shown that CD4+ T cells respond differently to phosphorylated and non-phosphorylated rabies virus antigens^9,10^ as well as lupus auto-antigens^11^ in lymphocyte proliferation assays. A few years later, Meyer *et al.*^12^ and Depontieu *et al.*^13,14^ directly identified natural MHC class II phospholigands via tandem mass spectrometry (MS2), demonstrating that these modified peptides can indeed be processed and presented. Depontieu *et al.*^14^ also showed that human CD4+ T cells can distinguish phosphorylated residues using cells raised against the melanoma antigen mutant B-Raf and its phosphorylated counterpart. Additionally, in the same study, the authors found that the phospho-MART-1 antigen, isolated from two melanoma cell lines and identified via MS2, was specifically recognized by CD4+ T cells in the context of its cognate HLA-DRB1 molecule.

While there is increasing evidence of the relevance of phosphorylated ligands in immune responses, their identification remains technically challenging. As stated above, the main source of data currently applied for characterizing and learning the rules of HLA antigen presentation is immunopeptidomics (reviewed for instance in ^15^). In short, a conventional immunopeptidomics pipeline consists of cell lysis, MHC-specific immunoprecipitation combined with liquid chromatography coupled to high-resolution tandem mass spectrometry followed by peptide identification by spectral annotation using software such as MaxQuant^16,17^, MSFragger^18^ and PEAKS^19^. While these pipelines have been greatly optimized and improved over the last decades, they have not been specifically adapted for stabilization and identification of post-translational modifications, making their application for the identification of MHC phospholigands particularly complicated. This, first and foremost, is due to the relatively low abundance of post-translationally modified peptides, and the fact that conventional enrichment techniques such as metal-affinity chromatography^20,21^ or titanium dioxide-based affinity chromatography^22^ are not commonly used in immunopeptidomics workflows which negatively impacts the phosphopeptide yield. Secondly, peptide ionization and fragmentation techniques are not specifically optimized for detecting MHC phospholigands. Thirdly, in addition to the already computationally demanding task of annotating MHC ligands from MS2 spectra, identifying the precise phosphorylation sites introduces an additional layer of complexity. This challenge arises from the existence of positional isomers: modified peptides that share the same molecular formula but differ in structure depending on which amino acid (serine, threonine, or tyrosine) is phosphorylated^23^.

These issues result in a very low data yield and quality when analyzing conventional MS-immunopeptidomics datasets for the presence of phosphorylated peptides. This is for instance illustrated in the earlier work by Solleder *et al*.^24^. In this work, the authors re-analyzed a larger set of MS-immunopeptidomics datasets using the MaxQuant spectral annotation software for the presence of phosphorylated ligands and found on average ∼100 peptides per sample (which is typically 10-50 times lower when compared to the identification of unmodified peptides^25^). Moreover, seeking to train a prediction model on these data, the authors concluded that close to half of these observed peptides may consist of co-eluted contaminants or wrongly identified peptides.

Given these limitations, we propose that novel workflows and advanced computational algorithms are needed for accurate identification and training of methods for prediction of MHC class II antigen presented phospholigands. To illustrate this, we defined a workflow refined for optimal PTM recovery, and used this to re-analyze the raw MS datasets from Solleder *et al.*^24^ Our results show that our workflow recovered a higher quantity and quality of HLA-II phospholigands, compared to that of the earlier work of Solleder *et al.* underlining the need for specialized immunopeptidome-specific workflows for accurate data acquisition. Given this, these datasets combined with others, likewise generated by re-analyzing MS-immunopeptidomics spectral datasets using our workflow, were used to train a ML model, termed NetMHCIIphosPan, which outperforms all prior state-of-the-art methods. This model fills a critical gap in existing prediction tools, such as NetMHCIIpan-4.3, which do not account for PTMs in their predictions, offering insights into the impact of phosphorylation on antigen presentation, which could be highly relevant in the context of autoimmune diseases, infections, and cancer.

## Materials and methods

### HLA-II MS Datasets

In this study, we employed HLA class II (HLA-II) phospholigands derived from various MS-based immunopeptidomics experiments. The PEAKS DeepNovo Peptidome, version 11.5 (Bioinformatics Solutions, Waterloo, Canada), hereafter referred to as PEAKS was used to search the raw MS data. In the final list of identified peptides, only peptides found by database search were included in the study. That is, peptides identified by homology search (peptides with mutations) or *de novo* sequencing were excluded.

Raw MS data from 20 single-allelic (SA) cell lines from Abelin *et al.*^26^ and 23 multi-allelic (MA) cell lines from Racle *et al.*^25^ were used in this search. Sequential HLA purification experiments were performed on 12 of the 23 multi-allelic cell lines, first with an HLA-DR monoclonal antibody (termed “Racle_DR” cell lines) and then with a pan-HLA-II antibody (termed “Racle” cell lines), yielding different subsets of ligands. Of the remaining, 10 multi-allelic cell lines were only processed with a pan-HLA-II antibody, and one was only processed with an HLA-DR monoclonal antibody. This resulted in a total of 35 subsets of ligands derived from multi-allelic cell lines.

Additionally, we included MS datasets from previous studies by Nilsson *et al.*^5,6^, Kaabinejadian *et al*.^4^, and van Balen and colleagues^27^. These datasets that were derived from either genetically engineered single-allelic (SA) or homozygous cell lines included 16 HLA-DQ, 14 HLA-DP, and 11 HLA-DR datasets from which HLA-class II was purified using locus-specific monoclonal antibodies, SPVL3, B7/21 and L243 respectively. This allowed us to expand our search to cover the HLA-DP and HLA-DQ as well.

The identified sequences were searched against the Swiss-Prot *Homo sapiens* (Taxon identifier 9606) database, including 20,395 entries, accessed on 12 May 2021 (https://www.uniprot.org). Precursor mass error tolerance of 10 ppm and fragment mass error tolerance of 0.02 Da were used. The enzyme was set to none and HCD (High-energy collisional dissociation) was selected as the fragmentation method. Variable post-translational modifications (PTM) including acetylation (N-term), deamidation, pyroglutamate formation, oxidation, sodium adducts, phosphorylation (STY), cysteinylation and citrullination were included in the database search. The maximum number of PTMs per peptide was set at 3 and the peptide length was restricted to 7 to 30 amino acids. Identified peptides were further filtered at a false discovery rate (FDR) of 5% using the PEAKS decoy-fusion algorithm, in line with earlier work^24^, to allow for an increased peptide yield. Default setting was used for the rest of the parameters including confident amino acid threshold and DeepNovo Score. For data acquired using the Triple-TOF Sciex 5600 instrument^4–6^, CID (Collision-induced dissociation) was selected as the fragmentation method, and precursor mass error tolerance was set at 30 ppm. Other parameters remained the same as above, and only peptides found by database search (DB Search) were included in the study.

In addition to the PEAKS analysis, the raw files from the Racle datasets were re-analyzed using FragPipe v23.1, incorporating MSFragger v4.3 and Philosopher v5.1.2^18,28,29^. MSFragger searches were performed using the same Swiss-Prot *Homo sapiens* database (accessed 12 May 2021) to ensure comparability with PEAKS output. Precursor mass tolerance was set to ±10 ppm and fragment mass tolerance to 0.02 Da. The enzyme specificity was set to “nonspecific”. Variable modifications were oxidation (M), phosphorylation (STY), cysteinylation (C) and protein N-terminal acetylation, with a maximum of 2 variable modifications per peptide. Peptide lengths were restricted to 7 to 30 amino acids. Identified peptides were filtered with sequential FDR at PSM- and protein-level 1% using Philosopher’s implementation of PeptideProphet. MS1-level peptide quantification was performed using IonQuant^30^, with default label-free quantification (LFQ) settings, including MaxLFQ, match-between-runs (MBR), and intensity normalization across runs. To ensure methodological consistency and enable a fair comparison with FragPipe, we conducted a second PEAKS search using the same search parameters.

All identified phosphopeptide sequences of length 12–21 amino acids are listed in **Supplemental Table S1**.

### Training Datasets

For all trained methods, negative data were generated as random natural phosphopeptides sampled for each phospholigand dataset, covering the same peptide length range as the latter, but with equal numbers of random peptides for each peptide length. More precisely, this number of random peptides was set to five times the number of ligands for the most prevalent peptide length. In this manner, the algorithm can learn the difference in peptide length preferences for ligands, as described before^31,32^. The random phosphopeptides were sampled from the human phosphoproteome downloaded from PhosphoSitePlus^33^, a comprehensive MS database of post-translational modifications (PTMs).

Several methods were trained on different datasets. Models M1, M2 (and its variants M2’ and M2’’) were trained on the original data from Solleder *et al.*^24^. Model M3 was trained on the same original MS data reanalyzed using the workflow defined here. Model M4-M6 were trained using an extended dataset, expanding the dataset from M3 with phospholigands obtained from earlier MS studies (Saghar_DR/DP/DQ^4–6^ and PvanBalen_DP^27^) reanalyzed using the workflow defined here. M4 was trained using only the extended phospholigand dataset. For M5 and M6, the training data was extended by unmodified MS eluted ligands derived from human cell lines or alleles, also from the NetMHCIIpan-4.3 training dataset. These datasets consist of single-allelic (SA) cell lines (101,449 peptides from 70 human cell lines covering 52 HLA-II alleles) and multi-allelic (MA) cell lines (44,956 peptides from 37 human cell lines covering 52 HLA-II alleles). For all models, a set of unmodified peptide binding affinity data (BA) extracted from the training dataset of the NetMHCIIpan-4.3 method (125,985 peptides covering 72 HLA alleles) was included in the training data. For the models M5 and M6, phospholigands and random natural phosphopeptides were upscaled (i.e. duplicated) 10 times when incorporating them into the training dataset.

### Data Partitioning

As described below, models were trained and evaluated using five-fold cross-validation. Before generating the five training data partitions, phosphosites (represented as “sty”; where “s” stands for pS, “t” for pT, and “y” for pY) were translated into their corresponding uppercase “STY” amino acids. Next, a common motif algorithm, inspired by the original Hobohm 1 algorithm, was used to cluster peptides sharing any identical 9-mer subsequences^34^. This ensures that nested peptides, ignoring phosphosites, are placed in the same partition. After partitioning, the phosphosites were back translated to “sty”. The final training dataset is available at https://services.healthtech.dtu.dk/services/NetMHCIIphosPan-1.0/, “Supplementary material” tab.

### Model Training

Models were trained using a five-fold cross-validation scheme, employing the NNAlign_MA algorithm^35^ The model architecture was set to 40 and 60 hidden neurons, and each architecture was trained with 10 random weight initializations per cross-validation fold resulting in a final ensemble consisting of 100 (5*10*2) network models. All models were trained using backpropagation with stochastic gradient descent, for 400 epochs, without early stopping, and a fixed learning rate of 0.05. Peptides were represented using sparse sequence encoding with an extended amino acid alphabet: ARNDCQEGHILKMFPSTWYVsty, where “sty” represents the phosphorylated amino acids pS, pT, and pY as described above. The motif length, or binding core, was set to 9 amino acids. Only SA data was included in the training for a burn-in period of 30 epochs, followed by training cycles including both SA and MA data. During the first 10 training iterations, only ILVMFYWRKsty amino acids were allowed at position 1 of the binding core. Models were trained including peptide flanking region (PFR) amino acid composition encoding, PFR length encoding, and peptide length encoding as described earlier^32^. If required, models were trained allowing for peptide inversion as described in ^6^.

### Independent Benchmark Datasets

External phospholigand datasets were gathered to benchmark the NetMHCIIphosPan method against other state-of-the-art tools, such as MixMHC2pred-1.3^24^. Two distinct class II immunopeptidomics datasets were selected, both acquired in the laboratory of Nicola Ternette. The first dataset consisted of unpublished data generated on Orbitrap Fusion Lumos Tribrid Mass Spectrometer from fresh frozen tissue surgically resected from 14 subjects diagnosed with Esophageal adenocarcinoma (EAC), tissue consisting of both tumor and normal adjacent Esophagus and Stomach. The second dataset consisted of previously published class II immunopeptidomes data from Monocyte-derived dendritic cells (MDDCs) generated on a Q-exactive HF-X Hybrid mass spectrometer from 5 HLA-DRB1-heterozygous healthy blood donors^36^. Here, each donor MDDC was pulsed with a recombinant SARS-CoV-2 S protein vaccine immunogen candidate. Both tissue and MDDC class II immunopeptidomes were generated using the anti-HLA-DR monoclonal antibody (L243). In the case of the “Esophageal tumor” dataset, the HLA typing only covers DRB1 alleles (and HLA-DQ alleles, though these were not employed for making predictions). For these datasets, DRB3/4/5 alleles were inferred with the HLAAssoc-1.0 tool^37^. Regarding the “Spike” dataset, full HLA typing is available for these samples.

#### Preparation of peptides from tissues and cells

Tissues were placed in a lysis buffer (0.5% (v/v) IGEPAL 630, 50 mM Tris pH 8.0, 150 mM NaCl, and 1 tablet complete Protease Inhibitor Cocktail EDTA-free (Roche) per 10 mL buffer) in bead beater tubes for soft tissues (Precellys Evolution, Bertin Technologies, France), bead beating was performed with 5 cycles at 7200 rpm for 20 s, separated by 20 s pauses followed by mixing for 30 min at 4°C to obtain complete lysis. Lysates were cleared by centrifugation initially at 500 *g* for 10 min and supernatants were removed and centrifuged at 21,000 *g* for 45 min at 4 °C to complete clarification. Peptide-HLA complexes were immunoprecipitated with L243 MAb cross-linked protein G sepharose beads at 4 °C overnight. The resin was washed consecutively in Tris buffer 50 mM, pH 8.0 containing 150 mM NaCl, 5 mM EDTA; 150 mM NaCl; 450 mM NaCl; no addition. Peptides were eluted twice with 500 μL 10% acetic acid and dried in a speed vac before clean-up by HPLC as described in ^36^.

#### LC-MS data acquisition

The Sars-CoV-2 dataset was generated as described^36^, using an Ultimate 3000 RSLC nano system (Thermo Scientific) to a Q-exactive HF-X Hybrid Mass Spectrometer (Thermo Fisher Scientific). For the EAC tissue, purified peptides were initially trapped, separated and spectra were acquired using an Ultimate 3000 RSLC nano system (Thermo Scientific) to an Orbitrap Fusion Tribrid Mass Spectrometer (Thermo Fisher Scientific) with a nano-electrospray ion source. Separation was achieved on a PepMap C18 EASY-Spray column (3 μm particle size, 75 μm x 50 cm) (Thermo Scientific) in buffer A (0.1% formic acid) using a linear gradient of 2%–35% buffer B (acetonitrile and 0.1% formic acid) at a flow rate of 300 nl/min over 60 min. The ion transfer tube temperature was 305 °C and full MS spectra was taken (300 to 1500 *m*/*z*) in the Orbitrap at 120,000 resolution with an automatic gain control (AGC) target of 40,000. Precursors were isolated using an isolation width of 1.2 amu and MS2 scans were acquired at maximal speed (120 ms with an AGC target of 300,000). Higher-energy collisional dissociation (HCD) with NCE of 28 was used to fragment peptides with a charge state of 2–4, for singly charged peptides NCE of 32 was applied with a lower priority in the decision tree. Fragments were analyzed in the Orbitrap at 30,000 resolution.

#### Phosphopeptide data analysis and benchmark dataset generation

Both datasets were analyzed with the PEAKS DeepNovo Peptidome 11.5 platform, applying the same search parameters and settings mentioned before for the training datasets (refer to the section entitled “HLA-II MS datasets”). The list of identified peptide sequences can be found in **Supplemental Table S2**.

As all external evaluation data originated from MS experiments, each dataset was enriched with random natural phosphopeptides in the same manner as for the training data. For the independent “Esophageal tumor” dataset, normal and tumor samples from each subject were combined into a single dataset. Only datasets with at least 10 positives were kept, resulting in a benchmark comprising 2058 phospholigands (available at https://services.healthtech.dtu.dk/services/NetMHCIIphosPan-1.0/, “Supplementary material” tab).

### Performance Evaluation

When assessing the performance of NetMHCIIpan-4.3 on our method’s training dataset and when evaluating the performance of NetMHCIIpan-4.3 and the method developed here on the external datasets, the following strategy was employed.

For NetMHCIIpan-4.3, which was trained exclusively on unmodified ligands, three “phospho-equivalent” datasets were generated: (1) phosphosites were replaced with glutamic acid (“E”) due to its similar charge, (2) phosphosites were substituted with amino acids of similar charge (pS → D, pT → E, pY → Y) (“DEY”), or (3) phosphosites were reverted to their unmodified forms (“STY”). Next, the overlap between the corresponding “phospho-equivalent” dataset and the NetMHCIIpan-4.3 training dataset was computed using the common motif algorithm^34^. For peptides with a 9-mer overlap with peptides placed in a given NetMHCIIpan-4.3 training partition, predictions were made using the neural network ensemble that was not trained on that specific partition (NetMHCIIpan-4.3 was trained using a five-fold cross-validation scheme, which implies that 20 out of the total 100 networks were used to make predictions). If no overlap was found, a random neural network ensemble (also consisting of 20 networks) was selected for predictions.

To evaluate our method, which was trained on both unmodified and phosphopeptides, phosphopeptides in the external datasets were converted from “sty” to “STY”. Then, the same evaluation procedure described before was followed: predictions were generated using the neural network ensemble trained on orthogonal data (as defined in the “converted” peptide space) to the evaluation data to prevent artificially inflated performance values.

### Performance Metrics

The model’s ability to distinguish ligands from random peptides was assessed using four performance metrics: the ROC-AUC (Area Under the Receiver Operating Characteristic Curve), the ROC-AUC 0.1 (ROC-AUC integrated up to a False Positive Rate (FPR) of 10%), the PR-AUC (Area Under the Precision-Recall Curve), and PPV. Note that ROC-AUC 0.1 values were not normalized using the McClish standardization; therefore, a random performance for this metric does not correspond to 0.5. The Positive Predictive Value (PPV) was computed as the proportion of true positives among the top N ranked peptides, where N corresponds to the number of ligands (positive instances) for a given HLA or cell line. In 5-fold cross-validation, the data was partitioned into five subsets. For each fold, models were trained on four-fifths of the data and evaluated on the remaining one-fifth, which served as the test set. This procedure was repeated five times, with each subset used once as the test fold. The resulting five subsets of test predictions, which together covered the entire dataset, were concatenated, and all performance metrics were computed on the combined set of “raw” prediction scores. For the independent benchmark, a similar approach was applied. For each peptide, models from a single training fold identified from a 9-mer overlap (with phosphosites being translated into their corresponding uppercase “STY” amino acids) to the training partitions were used for the predictions. In case no overlap was found, models from random training fold were used. Next, metrics were similarly calculated using “raw” prediction scores generated for each HLA allele or cell line. All metrics were reported only for datasets or cell lines containing at least 10 positive instances.

### Statistical Rationale

The corresponding statistical tests employed to compare model performances or other results are described in detail in the legend of each of the figures. In all cases, p-values less than 0.05 are considered to be statistically significant. In **Fig. 8**, pairwise comparisons of model performances were made using one-tailed binomial tests. For each comparison, 10,000 re-samples of the entire predicted benchmark dataset were generated, allowing for replacement (each subsample should contain at least 1% of the total original positive instances). Next, the p-value for one method outperforming the other was estimated as the fraction of re-sampled datasets where the other method had superior performance.

### Data Visualization

Sequence logos were made using the R “ggseqlog” package developed by Wagih *et al.*^38^. All the remaining plots in this study were made in R using the “ggplot2” package.

## Results

This work introduces NetMHCIIphosPan, a ML-based predictor of HLA class II presentation of phosphorylated peptides. The Results section is organized to reflect the progressive development of this tool. We begin by defining a workflow refined for optimal PTM recovery and show how this workflow results in higher MHC-II phosphopeptidome data quantity and quality compared to that reported in earlier studies^24^. We next train predictions models on the generated data and further investigate its consistency and quality. Further, we explore strategies to refine our method, including the incorporation of unmodified ligand data and support for inverted binding mode. Finally, we benchmark the final method against state-of-the-art tools, both in cross-validation and in an independent benchmark to demonstrate its robustness and improved performance.

### Data overview: improved phosphopeptide identification

As a starting point, we extracted the original HLA-II phosphorylated ligand dataset employed in Solleder *et al.*^24^. In this work, the authors re-processed high-quality MS samples from Abelin *et al.*^26^ and Racle *et al.*^25^ using oxidation (M), N-terminal acetylation and phosphorylation (STY) as PTM in their spectral searches with the MaxQuant software (version 1.5.5.1), and from these HLA-II phospholigandome data characterized the binding motifs for more than 30 HLA-II molecules. A main challenge, described in the work by Solleder *et al.*, concerning this HLA-II phospholigandome dataset, is the presence of co-eluted contaminants or wrongly identified peptides resulting in approximately half of the data being discarded by the authors.

We speculated that a substantial source of the seeming low quality of data generated in the study by Solleder *et al.* stems rather old version of the MaxQuant software used in the study and hypothesized that the issue could at least in part be corrected by using a more recent method for MS spectral annotation. Many such novel tools exist, and we here do not intend to suggest an optimal choice. Rather, we simply picked one such tool, PEAKS, for the task given our in-house expertise and experience with its usage. Based on a series of pilot studies, we define a workflow tailored to discover PTM modified peptides with properties aligning with MHC class II antigen presentation (peptide length distribution and MHC binding potential as defined from models trained on unmodified peptides). Using this workflow, we next re-process the raw MS samples from the Solleder *et al.* study. Further, we included other MS samples in our searches (Saghar_DR/DP/DQ^4–6^ and PvanBalen_DP^27^) to expand the allele coverage to better characterize HLA-DP and DQ binding motifs (refer to Materials and Methods for more details on datasets). Using this approach, we were able to increase peptide numbers and HLA coverage of the data in comparison to the original publication. **Fig. 1** provides an overview of the datasets re-processed with MaxQuant by Solleder *et al.* and the same datasets re-analyzed with PEAKS (refer to **Supplemental Fig. S1A-C** for the statistics over the full PEAKS datasets). Overall, the MaxQuant and PEAKS datasets display similar peptide length distributions preserving the peak at 15 amino acids long (**Fig. 1A**), although the former is shifted towards shorter peptide lengths. The peptide length distribution of unmodified HLA-II ligands falls between the distributions of these two datasets. The distribution of phosphosites in the two datasets is pS (55%/50%), pT (34%/36%), and pY (11%/14%) (for MaxQuant/PEAKS datasets, **Fig. 1B**), in line with the proportions of observed phosphosites in human proteins reported in PhosphoSitePlus^39^ (pS: 58%; pT: 25%; pY: 17%). Most of the phosphorylated ligands (85%/88%) contain only one phosphosite, while a small fraction contain two (13%/11%), and almost none contain three (2%/1%) phosphosites (for MaxQuant/PEAKS datasets, **Fig. 1C**). PEAKS recovers 33%, 246%, and 118% more phosphopeptides for the Abelin, Racle (data generated with pan class II antibody in the immunoprecipitation), and Racle_DR (data generated with a DR-specific class II antibody in the immunoprecipitation) datasets, respectively, in comparison to the original MaxQuant analysis **(Fig. 1D)**. Considering these three datasets re-analyzed in common, Abelin, Racle and Racle_DR, the overlap between phosphopeptides identified by PEAKS and MaxQuant was limited. In the Abelin dataset, only 15 phosphopeptides were identified by both tools, corresponding to 2.0% of MaxQuant and 1.0% of PEAKS identifications. For the combined Racle and Racle_DR datasets, 499 overlapping peptides were observed, representing 15.1% of MaxQuant and 4.8% of PEAKS identifications. Across these three datasets, only 3.8% of the total unique identified phosphopeptides overlap between the two approaches. Taking all datasets into account 19,019 phosphopeptides were identified with the PEAKS software, while 4,632 phosphopeptides were identified with MaxQuant (including only 12 to 21-mers in both cases).

**Fig. 1.**
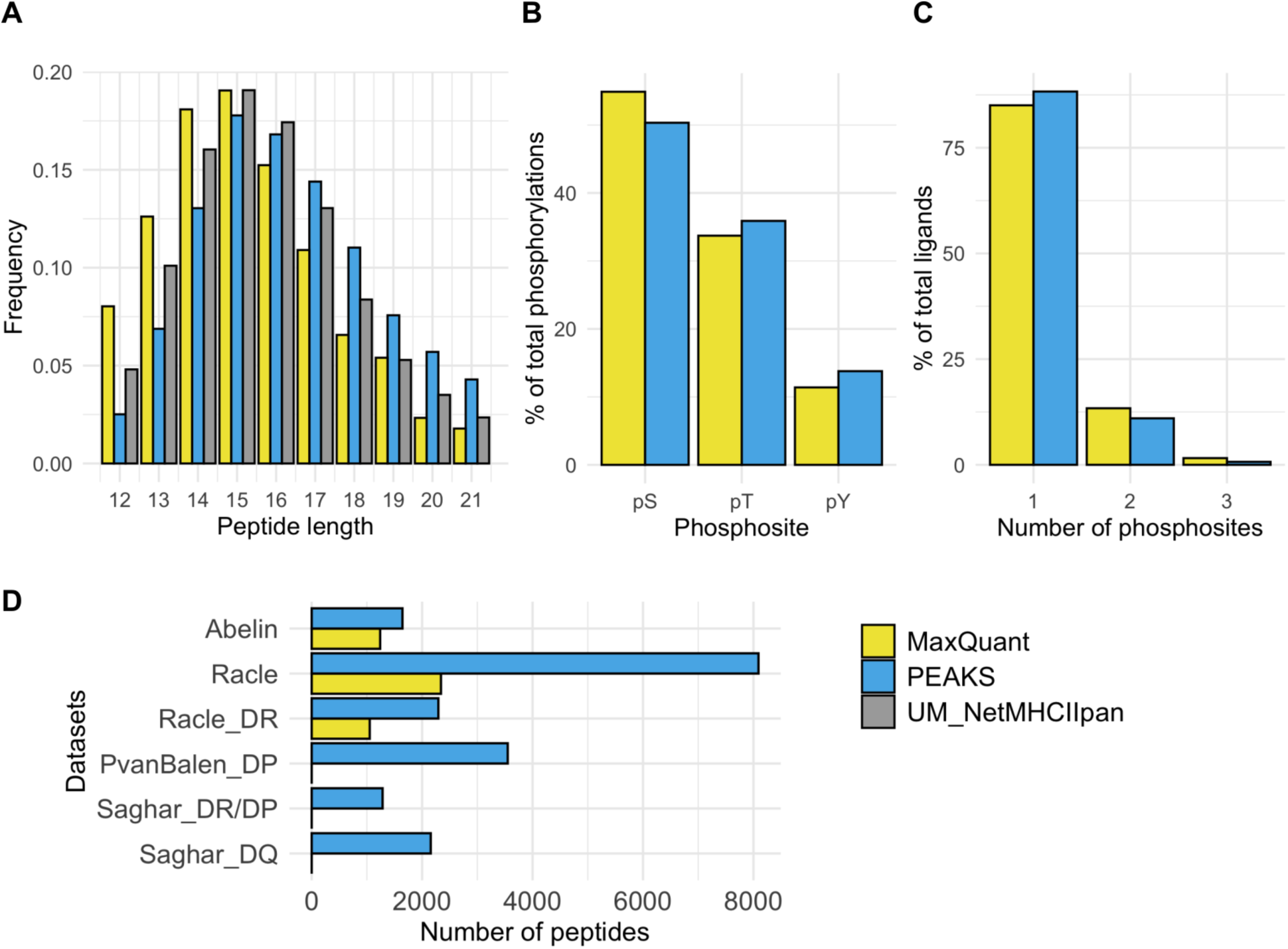
Overview of the phospholigand datasets identified by MaxQuant and PEAKS software. (**A**) Peptide length distribution. In grey, the distribution of unmodified HLA-II ligands lengths from the training dataset of the NetMHCIIpan-4.3 method (“UM_NetMHCIIpan”). (**B**) Phosphorylation type prevalence, considering all phosphosites found in modified peptides. (**C**) Distribution of the number of phosphosites per modified peptide. (**D**) Number of peptides identified in each of the analyzed datasets. Note that the PvanBalen and Saghar datasets were not analyzed using MaxQuant, and MaxQuant values are hence not available for these datasets in **D**.

We acknowledge that the comparison here is limited by the somewhat outdated version of MaxQuant used in the earlier study. We will return to this later in the manuscript, where we include a study using a third state-of-the-art algorithm for immunopeptidome identifications, and here demonstrate a high consistency to the PEAKS results. In summary, these results suggest that using the workflow defined here results in a major increase in peptide yields of identified phospholigands compared to that of earlier workflows.

### Enhanced predictive models for phospholigand binding

Next, to investigate the quality of the datasets and assess to what degree each could lead to an understanding of the HLA-II phospholigandome, models were trained either with the original data from the Solleder *et al.* study (Abelin, Racle and Racle_DR dataset re-analyzed using MaxQuant) or with data from the same EL datasets re-analyzed using the workflow defined here.

All developed models were trained using the NNAlign_MA machine-learning framework^35^. In short, the NNAlign_MA algorithm enables simultaneous training and motif deconvolution of multi-allelic (MA) data, i.e. MS eluted ligands from cell lines expressing multiple HLA-II alleles. This aspect is crucial for this work since a substantial amount of the phosphorylated HLA-II ligands employed in the training of these methods originate from multi-allelic data, which implies they have no pre-determined HLA-II allelic restriction. Indeed, this becomes even more relevant for this study as the HLA-II binding preferences of phosphorylated peptides have been very scarcely characterized and may differ from the HLA-II binding motifs described for their unmodified counterparts.

Models were trained using a five-fold cross-validation scheme. To prepare the training datasets, random negatives were sampled from the human phosphoproteome deposited in PhosphoSitePlus^33^, and peptides were grouped using a common motif algorithm, ensuring no 9-mer overlap between partitions^34^ (for details on Training Datasets, Data Partitioning or Model Training refer to Materials and Methods). Details related to the number of phospholigands (positive instances), random phosphopeptides (negatives) and the number of cell lines included in the training dataset of these models, are given in **Supplemental Table S3**. For each fold, the model was trained on four-fifths of the data and evaluated on the remaining one-fifth. This process was repeated five times, resulting in five sets of test set predictions that, together, covered the entire dataset. These predictions were concatenated, and metrics were computed on the combined set of prediction scores. We report four complementary performance metrics: ROC-AUC, ROC-AUC 0.1 which corresponds to ROC-AUC integrated up to a 10% FPR, Precision-Recall AUC (PR-AUC), and Positive Predictive Value (PPV) computed over the top N predictions, where N corresponds to the number of HLA-II ligands for each dataset. ROC-AUC 0.1 focuses on the high-specificity region of the ROC curve, which is critical when minimizing false positives, while PPV directly quantifies the model’s ability to prioritize true HLA-II ligands among top-ranked predictions, where experimental validation efforts are concentrated. Note, that the ROC-AUC 0.1 values reported are not normalized and that a random performance for this metric here corresponds to 0.05. Finally, it is relevant to mention that the NNAlign_MA framework requires both eluted ligand (EL) and binding affinity (BA) data to perform training. Therefore, in this case, the phospholigand training datasets were combined with more than 125,000 unmodified peptides measured in BA assays from the training dataset of the NetMHCIIpan-4.3 method.

**Fig. 2** illustrates the five-fold cross-validation performances of three methods, two of them, M1 and M2, trained with the MaxQuant datasets, and one trained with the PEAKS datasets (M3). For “MaxQuant_w_filter” M2 method, we applied the same filtering strategy as Solleder *et al.* using MixMHC2pred-2.0^40^ predictions replacing phosphosites by glutamic acid to filter the data and discard potential contaminants or misidentified peptides (%rank > 10). **Supplemental Fig. S2** expands this analysis including the performances of two other methods, trained on the MaxQuant datasets and filtered according to NetMHCIIpan-4.3 predictions.

**Fig. 2.**
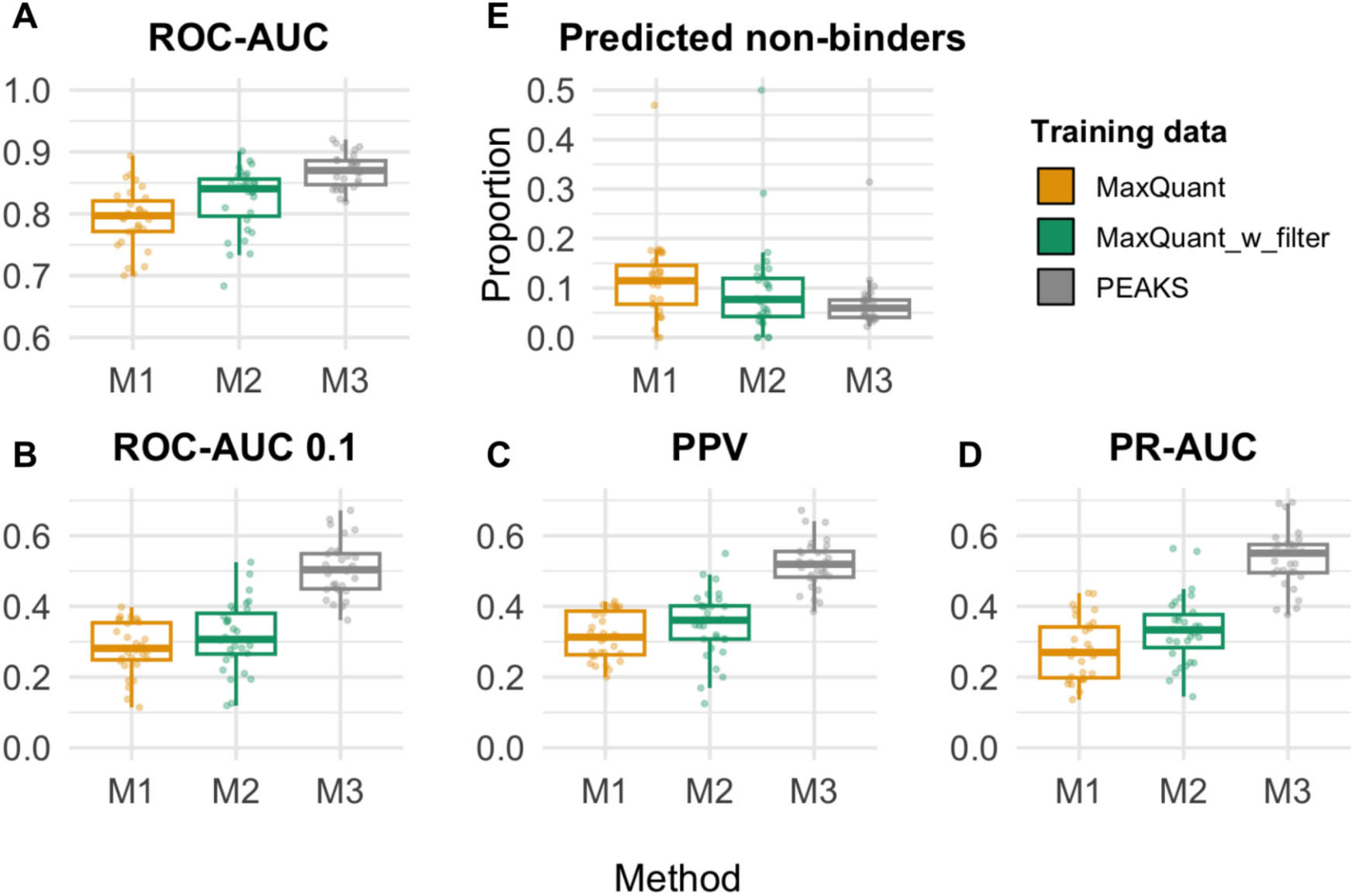
Five-fold cross-validation performance of the methods trained with phospholigands identified by MaxQuant and PEAKS. **M1** refers to the method trained and evaluated on the complete MaxQuant dataset, **M2** refers to the method trained and evaluated on the MaxQuant dataset, pre-filtered according to MixMHC2pred-2.0 predictions (%rank≤10), and **M3** refers to the method trained and evaluated on the complete PEAKS dataset. Performance metrics included are (**A**) ROC-AUC, (**B**) ROC-AUC 0.1, (**C**) PPV and (**D**) PR-AUC. ROC-AUC refers to the area under the Receiver Operating Characteristic curve; ROC-AUC 0.1 is the partial AUC integrated up to a FPR of 10%; PR-AUC is the area under the Precision-Recall curve; and PPV (Positive Predictive Value) is calculated as described in Materials and Methods. Performance is evaluated in a per-dataset manner from the concatenated phosphopeptide test sets. (**E**) Proportion of predicted non-binders (%rank>20) according to each of the methods. The center line inside the box indicates the median value of the plotted metric and the box covers the interquartile range. The whiskers extend to, at most, 1.5-fold of the interquartile range. The individual data points in the jitter plots correspond to different cell lines. Only cell lines with 10 or more positive instances for all trained methods were included in the analysis (N=31 cell lines). Y-axis ranges differ between panels **A** and **B**-**D**.

We notice that the different methods are trained and evaluated on datasets of varying sizes, which complicates direct comparison. Nonetheless, the results presented in **Fig. 2** and **Supplemental Fig. S2** show that, although filtering of potential noise improved the performance of the models trained and evaluated on the MaxQuant datasets, the model trained on the full, unfiltered PEAKS datasets consistently exhibits a superior performance. This enhanced predictive power is also reflected in the overall lower proportion of predicted non-binders (phosphopeptides with %rank > 20) within the PEAKS datasets, in relation to the MaxQuant and MaxQuant_w_filter datasets (refer to **Fig. 2E**). This robustness further contributes to the superior performance even in cell lines with very few data points. For instance, the M3 model achieves an average ROC-AUC 0.1 of 0.56 on the 10 datasets with the fewest positive examples (average of 130 positive data points per dataset), while the M1 model achieves an average ROC-AUC0.1 of only 0.25 on the corresponding 10 datasets with the most positive examples (average of 183 positive data points per dataset) (see **Supplemental Table S4**). This result suggests that the superior performance of the PEAKS-trained model is attributed not only to the increased data quantity but also a higher data quality.

### Improved motif consistency

We next evaluated the similarity between the HLA-II binding motifs of unmodified and phospholigands, for each of the trained models. For this, we computed position frequency matrices (PSFMs) based on the HLA restriction and predicted 9-mer binding cores obtained in five-fold cross-validation. Here, peptides with a predicted %rank score greater than 20% were considered potential HLA irrelevant co-immunoprecipitation outliers and were excluded from this motif comparison analysis. Likewise, phosphopeptides lacking phosphosites within the predicted binding core were also discarded. For more details on phosphosite localization within the binding core or peptide flanking regions see subsequent analyses and refer to **Supplemental Fig. S3**. Finally, only HLA-II molecules characterized with more than 30 ligands were included in the analysis. For unmodified peptides, binding cores from the cross-validation predictions from the training of the latest NetMHCIIpan-4.3 method were employed. Subsequently, PCCs (Pearson Correlation Coefficient) were computed between the flattened PSFMs corresponding to HLA-II phospholigand and unmodified ligand binding motifs. It is relevant to note that to enable this comparison of the binding preferences between modified and unmodified peptides, only the 20 unmodified amino acids were included in the estimates of the position frequency matrices (PSFMs).

**Fig. 3A** shows the distribution of PCCs for each of the trained models. An extension of this analysis can be found in **Supplemental Fig. S4**, which includes the results for two other models trained with the MaxQuant datasets and filtered according to NetMHCIIpan-4.3 predictions. In line with the results in **Fig. 2**, the model M1 displays the lowest concordance between HLA-II phospholigand and unmodified ligand binding motifs, followed by the model M2. Finally, the model M3 presents the highest concordance, with a median PCC of 0.82. For this model, the DQ molecules (DQA1*03:01-DQB1*03:02, DQA1*01:01-DQB1*05:01 and DQA1*03:03-DQB1*03:01) presented the lowest PCC values (PCCs < 0.3), sharing a low number of phospholigands assigned during training (N < 70 for all three molecules). After the DQ molecules, the next lowest correlations were observed for three DP molecules (PCCs < 0.6). Together, these results suggest that this model remains challenged in the characterization of the HLA-DP and DQ phospholigand space.

**Fig. 3.**
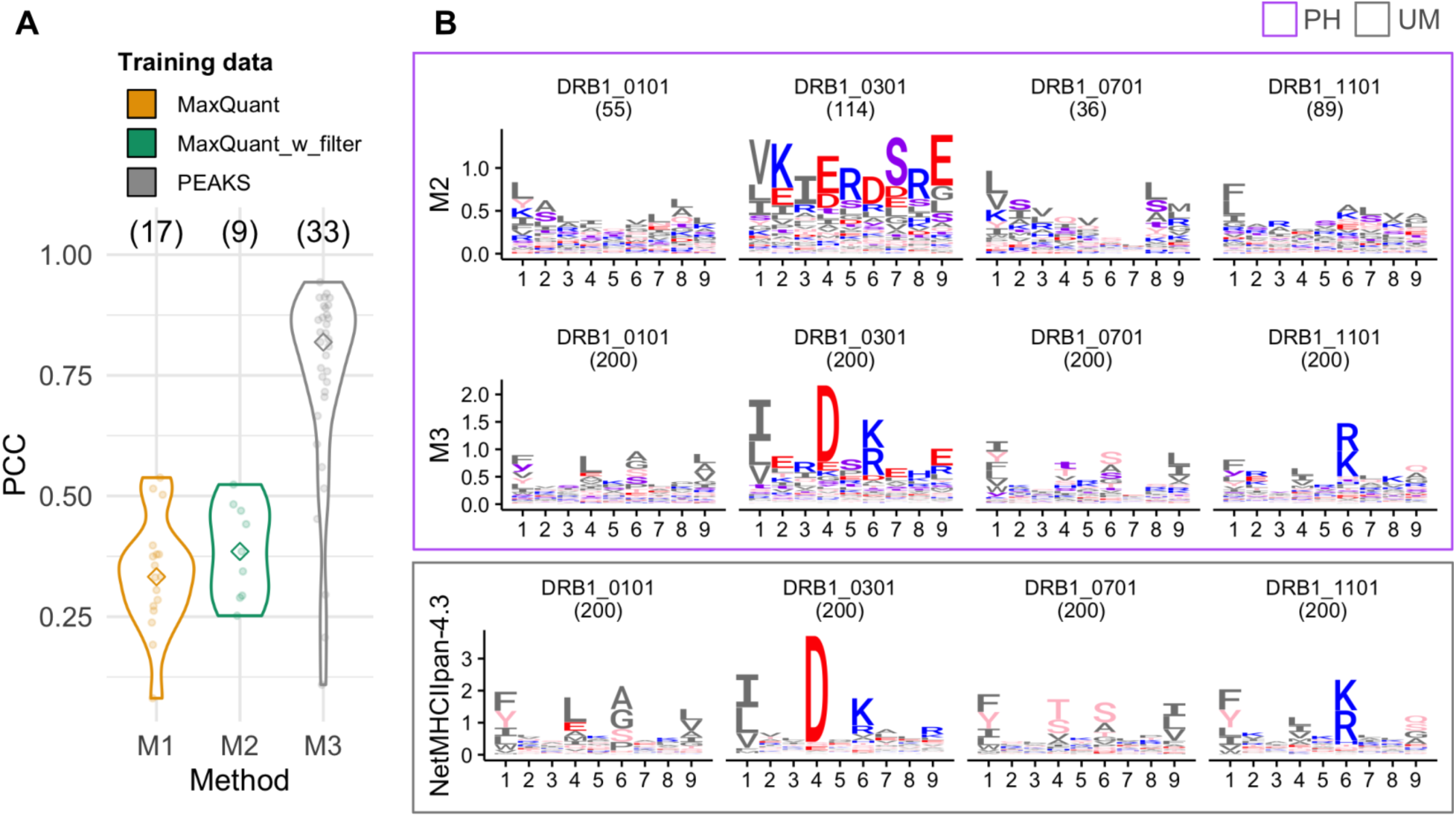
Concordance between unmodified and phospholigand binding motifs for HLA-II molecules with 30 or more peptides assigned during cross-validation. (**A**) PSFMs (position-specific frequency matrices) were computed to represent HLA-II binding motifs for unmodified and phosphopeptides, based on cross-validation predictions of the different methods. The three methods included are **M1**, **M2** and **M3** described in Fig. 2. In the case of phosphopeptides, phosphosites were ignored and frequencies per position were renormalized to sum up to 1. For unmodified peptides, predictions from NetMHCIIpan-4.3 were employed. To compare binding motifs for a given allele, PCCs (Pearson Correlation Coefficient) were calculated between flattened unmodified and phosphopeptide PSFMs. The violin plots show the distribution of PCCs for each of the trained methods (each data point corresponds to a different HLA-II molecule, the diamond indicates the median PCC, and, above each plot, the analyzed number of HLA-II molecules is noted). (**B**) Sequence logos for a subset of four selected DRB1 alleles, generated from cross-validation predictions as in (**A**). The first row corresponds to logos illustrating phospholigand (“PH”, outlined in violet) binding motifs for 4 HLA-II molecules according to the **M2** method, while the second row shows the same for the **M3** method. The third row shows logos representing unmodified (“UM”, outlined in grey) HLA-II ligand binding motifs according to NetMHCIIpan-4.3. Each logo displays in parenthesis the number of peptides employed for its generation. In the case of the **M3** and NetMHCIIpan-4.3 methods, logos were constructed by randomly subsampling 200 peptides per allele from the cross-validated test set predictions. Amino acid color coding: basic in blue (RHK), acidic in red (DE), polar in pink (STNQYC), hydrophobic in dark grey (GPLIVAMFW), and phosphosites in violet (sty).

Next, **Fig. 3B** illustrates sequence logos representations of binding motifs for four selected DRB1 alleles. Shannon logos were constructed employing the calculated PSFMs of models M2 and M3. Logos built on the M2 method appear fuzzy for all alleles, with barely defined anchor positions (typically positions 1, 4, 6, and 9) except for position 1. In contrast, logos corresponding to the M3 model appear sharper, with most anchor positions well-defined and in close concordance with the anchor positions observed in unmodified peptide logos of NetMHCIIpan-4.3.

In summary, the results from **Fig. 2** and **3** extend the conclusions from **Fig. 1**, detailing that not only does the novel pipeline proposed here result in a higher peptide yield compared to that of the earlier studies, but also suggest that the data is of a higher quality. Further, the results demonstrate that the M3 model, in addition to having higher predictive performance, also can infer more coherent HLA-II binding preferences from the training data, congruent with the existing knowledge on how unmodified ligands bind HLA-II molecules.

### Integrating unmodified and phospholigand data

In light of these results, and in line with conclusions from earlier work on HLA class I^41^, we next investigated whether the predictive performance of the M3 model could be improved by expanding the training set to include unmodified ligands. To achieve this aim, we constructed a new training dataset by combining phospholigands with over 140,000 unmodified HLA-II ligands, extracted as a subset of the NetMHCIIpan-4.3 training dataset. This combined set also included phospholigands derived from the additional van Balen and Saghar datasets described in **Fig. 1**. To balance out the number of unmodified and phospholigands included in the training dataset, phospholigands were upscaled by a factor of 10 (refer to Methods for more details). Furthermore, we also allowed for the inverted peptide binding described earlier for unmodified peptides for a subset of HLA-DP molecules^6,40,42^. **Supplemental Table S3** details the number of phospholigands, negative phosphopeptides, and cell lines used for model training.

**Fig. 4A-D** shows the five-fold cross-validation performance of the model M4, trained on phospholigands only, the model M5, trained on phospholigands and unmodified ligands, and the model M6, trained on the same data but including the option for the inverted binding motif. In terms of the ROC-AUC, ROC-AUC 0.1, PPV and PR-AUC metrics, the M5 and M6 methods demonstrated close to equal performance and significantly outperformed the M4 method (one-tailed binomial tests, for the ROC-AUC all P<0.05, and ROC-AUC 0.1, PPV and PR-AUC all P<0.0001). Next, we repeated the analysis to evaluate the similarity between unmodified ligand and phospholigand binding motifs for the studied methods, also based on model cross-validation predictions. As before, only HLA-II molecules with more than 30 assigned ligands were included. The results of this analysis are presented in **Fig. 4E**, comparing the three methods across different loci. For HLA-DR, the distribution of PCC values is highly similar for all three methods. For the HLA-DQ alleles, there is a marked difference between the PCC distributions of the two models trained including unmodified data to the model trained without. Here, the M5 and M6 (including inversions) methods both display a median PCC > 0.77, while for the M4 method this value is 0.60. Finally, for HLA-DP, the M6 model shows higher PCC values compared to the two other methods, with a median PCC of 0.91. Inspecting sequence logos representing phospholigand binding motifs for selected HLA-DP and HLA-DQ molecules (refer to **Supplemental Fig. S5**), it becomes clear that the motifs of the M5 and M6 methods mimic more closely the binding preferences found in the unmodified ligands (in this case represented by the NetMHCIIpan-4.3 logos). In particular, the sequence logos for the two exemplary HLA-DP molecules illustrate that the concordance between unmodified and phosphopeptide binding motifs is maximized for the M6 model, which allows for the inverted peptide binding mode, resulting in the consistent placement of “K” at P1 instead of P9. **Supplemental Fig. S6A** shows that the presence of the inverted binding mode for phospholigands follows the same rules as earlier described for unmodified ligands. That is, this binding mode is almost solely present for HLA-DP molecules, and mainly for molecules sharing the HLA-DPA1*02:02/01 alleles (see **Supplemental Fig. S6B**).

**Fig. 4.**
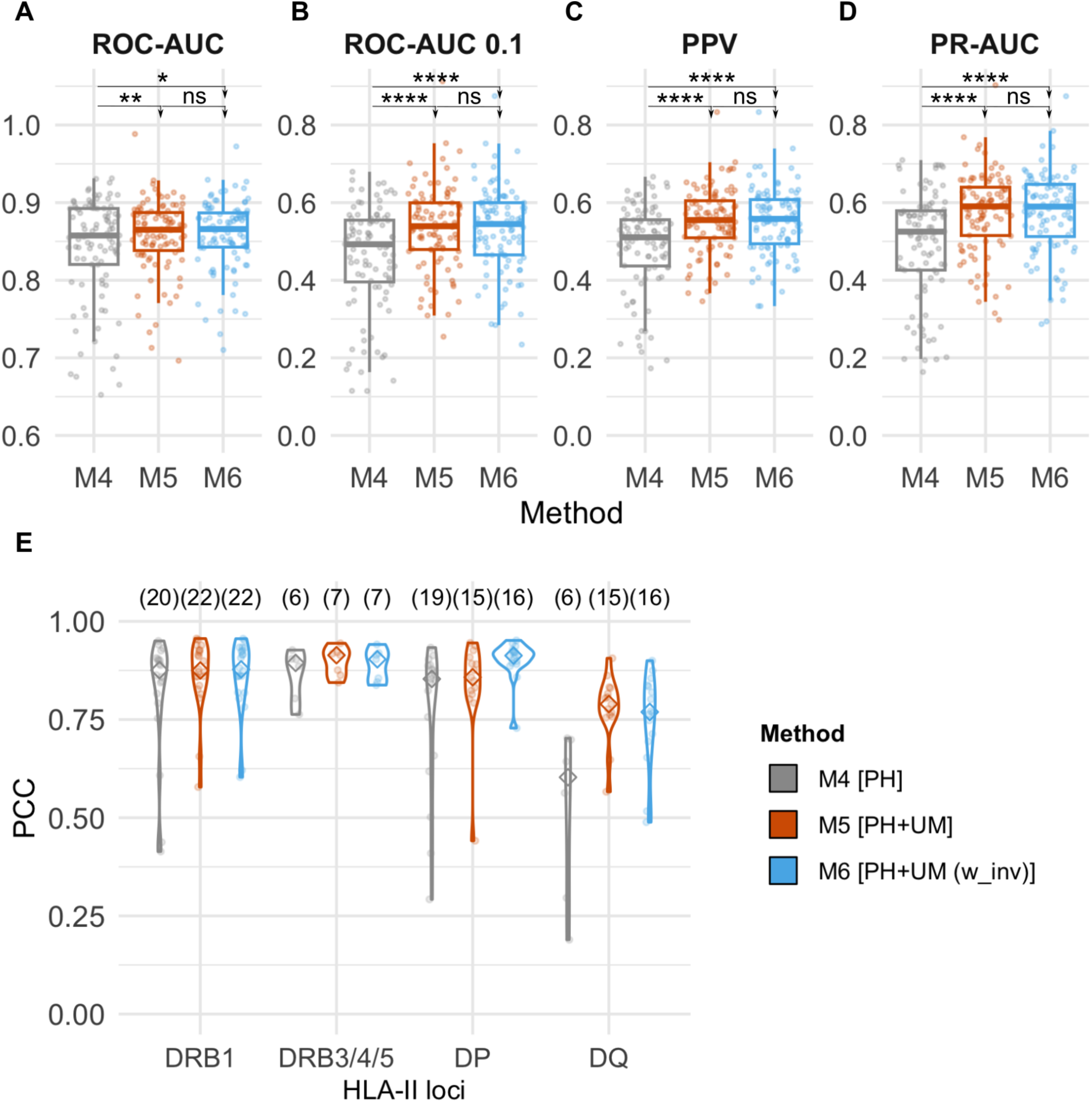
Improvement of the baseline method trained with phospholigands by adding unmodified ligands to the training dataset and allowing for peptide inversion. Five-fold cross-validation performance of the **M4** method, trained only with phospholigands (“PH”), the **M5** method, trained with both phospholigands and unmodified ligands (“PH+UM”), and the same method, **M6**, trained with peptide inversion (“PH+UM (w_inv)”; N=96 cell lines). Performance metrics included are (**A**) ROC-AUC, (**B**) ROC-AUC0.1, (**C**) PPV and (**D**) PR-AUC. ROC-AUC refers to the area under the Receiver Operating Characteristic curve; ROC-AUC 0.1 is the partial AUC integrated up to a FPR of 10%; PR-AUC is the area under the Precision-Recall curve; and PPV (Positive Predictive Value) is calculated as described in Materials and Methods. Performance evaluation, computed over the concatenated phosphopeptide test sets, and plots were made as in Fig. 2. One-tailed binomial tests were performed to compare the performance of the methods, with arrowheads indicating the direction of the tests (* P<0.05, ** P<0.01, and **** P<0.0001; ns, not significant). Y-axis ranges differ between panels **A** and **B-D**. (**E**) Concordance between unmodified and phospholigand binding motifs, based on model cross-validation predictions, for HLA-II molecules with 30 or more peptides assigned discriminated by loci. (plots were made as in Fig. 3A; above each plot, the number of analyzed HLA-II molecules is indicated).

In summary, these results demonstrate that the inclusion of the unmodified ligands and the option for the inverted binding mode have aided the deconvolution of complex multi-allelic phospholigand datasets, especially enhancing the characterization of HLA-DP and HLA-DQ binding motifs, which contributes to the overall moderate improvement in model performance. From now onwards, the M6 method will be referred to as NetMHCIIphosPan, our method of choice.

### Benchmarking against the state-of-the-art method: NetMHCIIpan-4.3

Next, we benchmarked the NetMHCIIphosPan method against its closest predecessor, NetMHCIIpan-4.3. To compare with the earlier developed method, solely trained on unmodified ligands, three “phospho-equivalent” training datasets were generated: (1) phosphosites were replaced with glutamic acid (“E”) due to its similar charge, (2) phosphosites were substituted with amino acids of similar charge (pS → D, pT → E, pY → Y) (“DEY”), or (3) phosphosites were reverted to their unmodified forms (“STY”).

**Fig. 5** shows that NetMHCIIphosPan significantly (all P<0.0001) surpassed NetMHCIIpan-4.3, evaluated over the different “phospho-equivalent” datasets across all metrics (for performance evaluation details refer to Methods section).

**Fig. 5.**
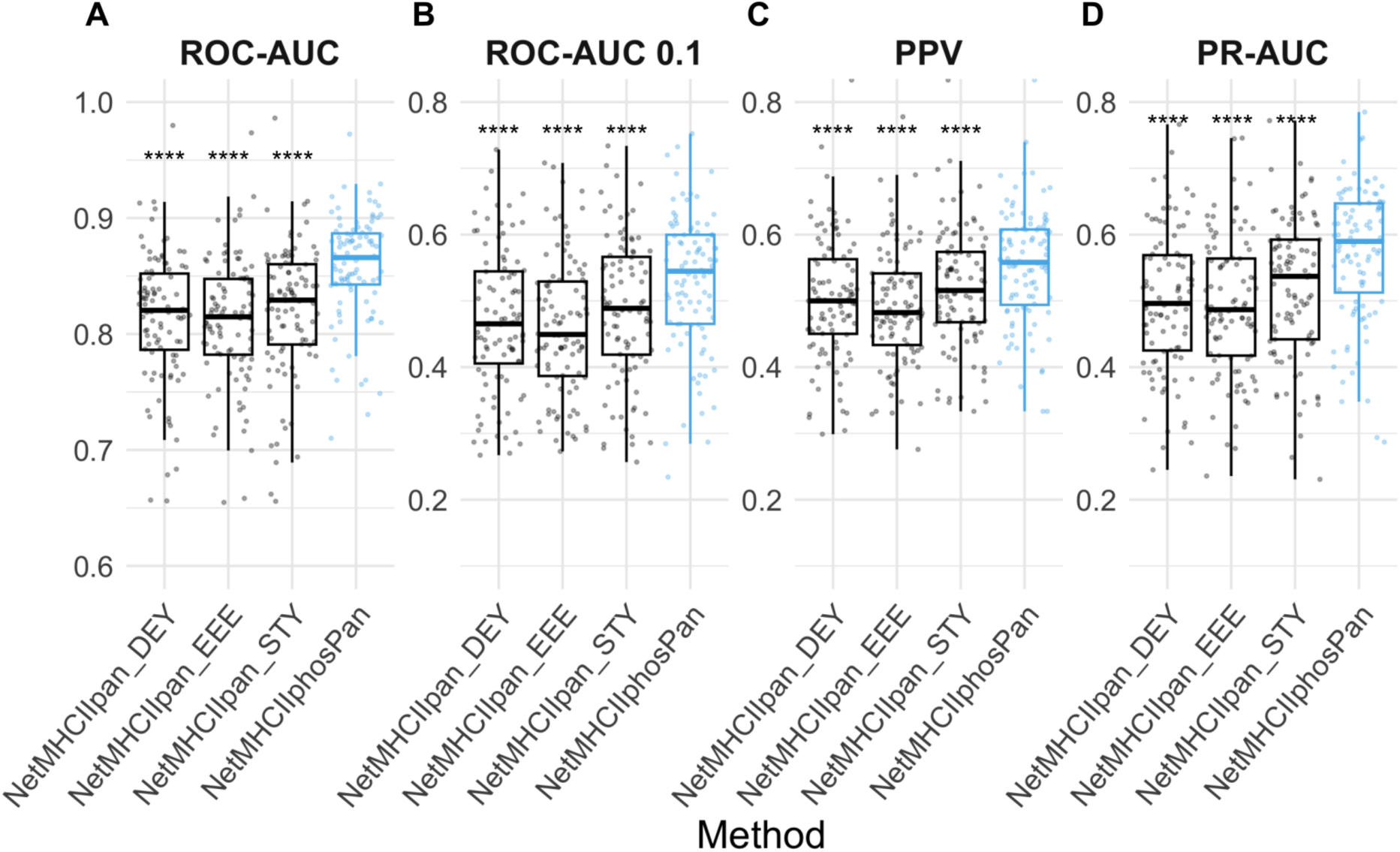
Five-fold cross-validation performance of NetMHCIIphosPan and NetMHCIIpan-4.3 on training dataset phospholigands. Performance metrics included are (**A**) ROC-AUC, (**B**) ROC-AUC0.1, (**C**) PPV and (**D**) PR-AUC. ROC-AUC refers to the area under the Receiver Operating Characteristic curve; ROC-AUC 0.1 is the partial AUC integrated up to a FPR of 10%; PR-AUC is the area under the Precision-Recall curve; and PPV (Positive Predictive Value) is calculated as described in Materials and Methods. Performance is evaluated in a per-dataset manner from the concatenated test sets (N=96 cell lines, for details see Materials and Methods). The center line inside the box indicates the median value of the plotted metric and the box covers the interquartile range. The whiskers extend to, at most, 1.5-fold of the interquartile range. The individual data points in the jitter plots correspond to different cell lines. One-tailed binomial tests were performed to compare the performance of the NetMHCIIphosPan method against the NetMHCIIpan-4.3 method (different “phospho-equivalent” datasets) (**** P<0.0001). Y-axis ranges differ between panels **A** and **B-D**.

### Concordance between unmodified and phosphorylated peptide binding motifs

Following, we assessed if the allele-specific binding motif preferences captured by NetMHCIIphosPan aligned with the motifs present in the training data. This analysis is essential to assess to what degree the model has learned the underlying motifs. Here, we compared logos for the 25 HLA-II molecules with the highest number of assigned phospholigands during training constructed either from the five-fold cross-validation NetMHCIIphosPan predictions (excluding phospholigands with predicted %rank > 5 and without phosphosites within the binding core in order to focus the analysis on peptides with very high antigen presentation likelihood only) or from the top 200 (0.1%) NetMHCIIphosPan predictions from a set of 200,000 random phosphopeptides (refer to NetMHCIIphosPan logos in **Supplemental Fig. S7**). As before, we evaluated the pairwise similarity between unmodified and phospholigand logos for each individual HLA-II molecule by computing the Pearson correlation coefficient (PCC) between their corresponding flattened PSFMs. **Supplemental Fig. S8** shows the distribution of PCCs, with a median value of 0.90, confirming that the motifs captured by the model to a high degree align with the one found in the phospholigands training datasets.

Subsequently, we created sequence logos and visually compared the phospholigand HLA-II binding motifs learned by our method with the corresponding NetMHCIIpan-4.3 motifs for unmodified ligands. To achieve this aim, we, like above, predicted two sets of 200,000 random phospho- and unmodified peptides each and selected the top 200 (0.1%) binders. **Fig. 6** presents sequence logos for ten alleles selected from the top 25 with the highest number of assigned phospholigands during NetMHCIIphosPan training. **Supplemental Fig. S7** extends this analysis to include all top 25 HLA-II molecules, as previously described. This analysis again confirms the high concordance between the binding motifs of phosphorylated and unmodified ligands. Focusing on the phospholigand motifs, the logos reveal that phosphosites (shown in violet) are distributed throughout the binding motifs, though they predominantly localize to canonical HLA-II anchor positions.

**Fig. 6.**
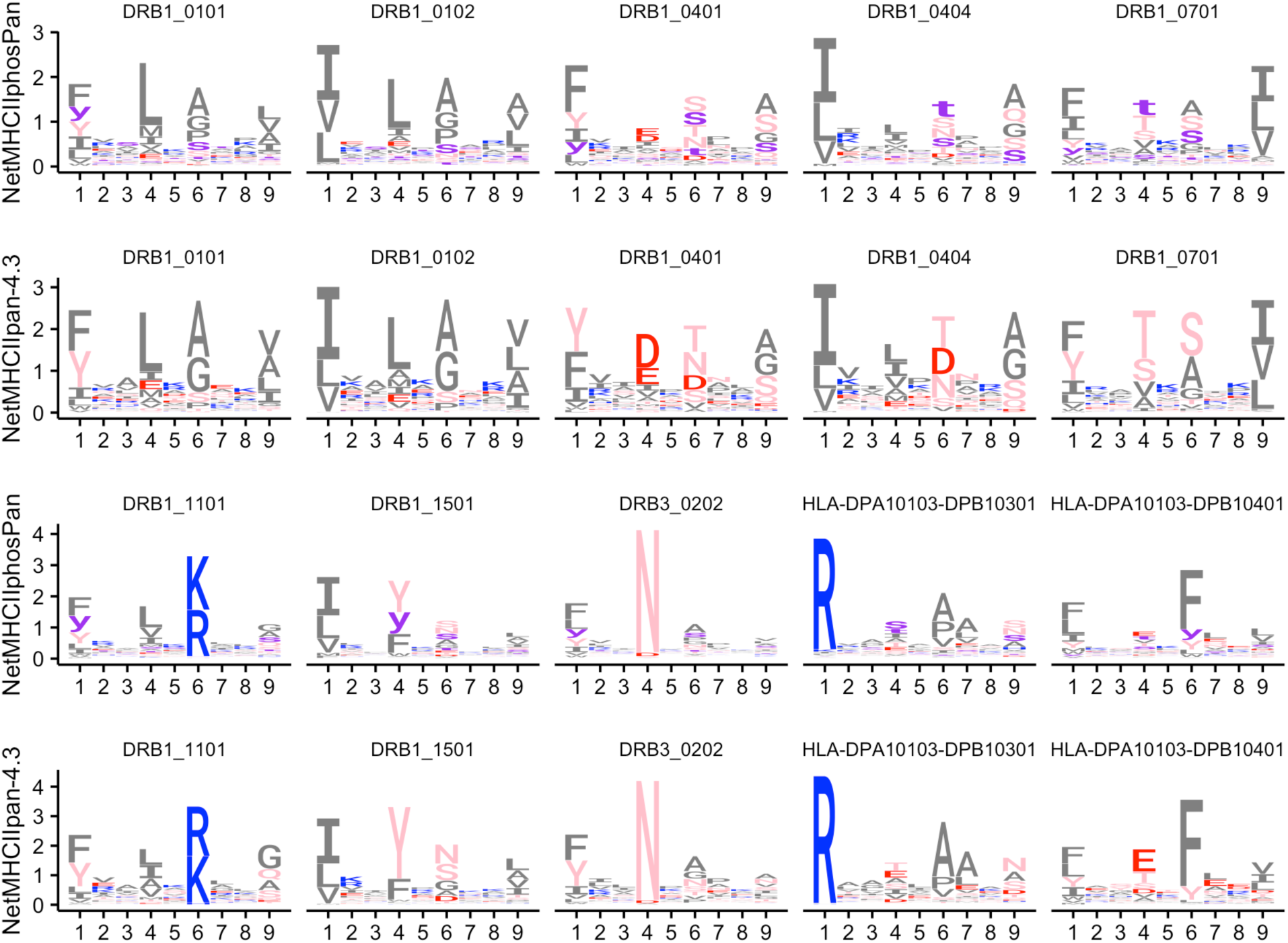
Sequence logo representations of HLA-II binding preferences for unmodified and phosphorylated peptides. Ten alleles were selected from among the 25 with the highest number of assigned phospholigands during NetMHCIIphosPan training. To construct the motifs two sets of 200,000 phospho- and unmodified peptides each were randomly sampled and predicted with NetMHCIIphosPan and NetMHCIIpan-4.3 respectively. The top 0.1% scoring peptides were used to construct the logos (all peptides had a predicted %rank≤5, and phosphopeptides without phosphosites in the binding core were excluded). Amino acid color coding: basic in blue (RHK), acidic in red (DE), polar in pink (STNQYC), hydrophobic in dark grey (GPLIVAMFW), and phosphosites in violet (sty).

### Insights into phosphosite localization preferences

Given this last observation, we next investigated in more detail if phosphosites shared a specific localization preference within the HLA-II ligands. This was evaluated first within the phospholigand with respect to the NetMHCIIphosPan predicted 9-mer binding core, and secondly within the predicted binding core with respect to the 9 possible binding positions. For the first analysis, we calculated the relative frequency of appearance of the phosphosite within the binding core or in its corresponding left or right peptide flanking regions (PFRs), averaged across peptides 13 to 17 amino acids long. This peptide length selection covers 69% of total phospholigands (refer to **Fig. 1A**) and was intentionally defined to be symmetric around 15-mers to compute an overall representative mean relative frequency. Taking natural random “non-binding” phosphopeptides (peptides with predicted %rank > 20) as a baseline, we observe (see **Supplemental Fig. S3**) that all three categories of phospholigands (%rank: [0,1], (1,5], (5,20]) behave similarly to this baseline, indicating that there is no particular preference with respect to the phosphosite localization between the binding core and PFRs.

Following this, we studied if any of the 9 binding core positions were enriched or depleted in phosphorylated amino acids. HLA-II binding predictions were computed for 200,000 random natural phosphopeptides for HLA molecules with 100 or more phospholigands assigned during model cross-validation (in total 50 HLAs). For each of the selected HLAs, the top 2.5% scoring predictions were taken as binders (top 5000 peptides per HLA), and an equal number of randomly sampled natural phosphopeptides with a predicted %rank > 20 were considered non-binders.

**Fig. 7** shows the log2 ratio between the relative frequency of binders vs. non-binders for each of the 9 binding core positions, discriminating by HLA loci/subtype and phosphorylation type. Overall, there are very limited consistent phosphosite enrichment/depletion signals. One exception being a few DRB1 alleles that are suggested to have enriched preference for pY at P1. In terms of depletion, most such observations appear to be located at the conventional class II anchor positions. Key examples of such marked HLA anchor position depletions include both pS and pT at P1 for almost all DRB1, DRB3/4/5, and HLA-DP molecules, and pY at P1 for most HLA-DP molecules, pY at P6 and P9 for DRB1 molecules, pY at P9 for HLA-DP and pY at P6 and P9 for HLA-DQ molecules.

**Fig. 7.**
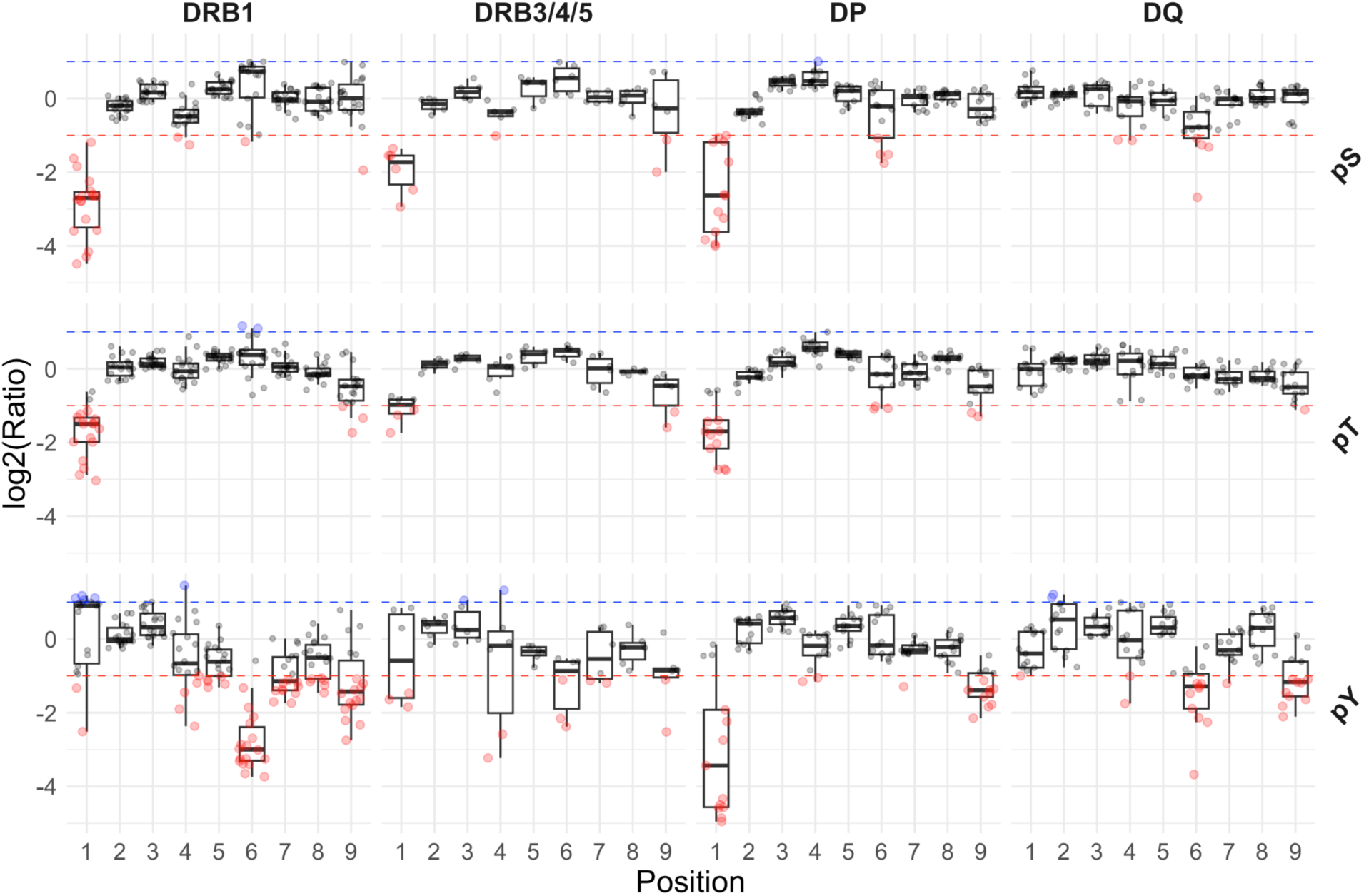
Phosphosite enrichment/depletion within the NetMHCIIphosPan predicted 9-mer binding core, discriminated by allele (columns) and phosphorylation type (rows). HLA-II binding predictions were computed for 200,000 random natural phosphopeptides for alleles with 100 or more phospholigands assigned during model cross-validation (in total 50). For each of the selected alleles, the top 2.5% scoring predictions were taken as binders, and an equal number of randomly sampled phosphopeptides with a %rank>20 were considered non-binders. The y-axis shows the log2 ratio between the relative frequency of binders vs. non-binders for each of the 9 binding core positions. Each data point in the plot corresponds to a different allele and, data points colored in blue indicate an enrichment of binders relative to non-binders (the blue dashed line corresponds to a log2 ratio of 1) whereas data points in red show the opposite, a depletion of binders relative to non-binders (the red dashed line corresponds to a log2 ratio of −1). **Supplemental Table S5** includes the computed log2(Ratio) for each binding core position, HLA-II molecule, and phosphorylation type.

Overall, these observations, together with the HLA binding motif analyses discussed earlier, suggest that phosphosite location is to a high degree aligned with the preferences described for unmodified S/T/Y residues. This is further supported by **Supplemental Fig. S9**, which compares the log₂ binder/non-binder ratios derived from random natural phosphopeptides with those obtained from random natural unmodified peptides across all nine core positions, stratified by HLA-II loci and residue type. When considering phosphosite depletion (log₂ ratio ≤ −1), a strong concordance with unmodified peptides is observed, with 92% (192/208) of depleted phosphosites matching depleted S/T/Y positions in unmodified peptides, of which 91% (175/192) are located at canonical HLA-II anchor positions (P1, P4, P6, and P9). These results indicate that phosphorylated S/T/Y residues are disfavored at certain anchor positions largely since their unmodified counterparts are themselves not preferred at these same positions. In contrast, among allele–position combinations showing phosphosite enrichment (log₂ ratio ≥ 1), only 58% (7/12; with no overlap observed for HLA-DP or HLA-DQ) coincide with core positions enriched for unmodified S/T/Y residues, and all such coincident cases occur at anchor positions. Together, these observations indicate that, within the current prediction framework, the impact of phosphorylation on HLA-II binding is predominantly constrained by the underlying enrichment/depletion patterns defined for the corresponding unmodified residues.

To further examine the impact of phosphorylation on peptide binding, we compared NetMHCIIphosPan cross-validation predictions for phospholigands in the training data and their unmodified counterparts (**Supplemental Fig. S10**). As in the earlier analyses, phospholigands with a predicted %rank > 20 were excluded as likely HLA-irrelevant co-immunoprecipitated contaminants. Among phosphopeptides that also appeared in unmodified form in the training dataset (10% of total), 95% retained the same predicted binding core, while only 5% were assigned an alternative one. For the remaining 90% without matched unmodified forms, 91% conserved their predicted binding core, and 9% exhibited a core shift. Thus, in the vast majority of phospholigands (91%), the predicted binding core remains conserved upon dephosphorylation, and for this subset, changes in binding likelihood were generally small, as indicated by the narrow distribution of delta %rank values centered around zero (see **Supplemental Fig. S10**). In contrast, phospholigands with a shifted binding core exhibited delta %rank values of larger magnitude, although still modest in most cases. In both groups, a prevalent trend toward increased %rank upon dephosphorylation was observed, reflected by the density shift toward positive values in the delta %rank distributions, indicating that removal of the phosphate group tends to reduce predicted binding likelihood. Focusing on the most relevant subset of phospholigands, those predicted as binders by our method (%rank ≤ 5), which account for 74% of the training dataset (14,046 out of 19,019), only 2.2% exceeded a %rank of 10, and just 0.37% surpassed 20 upon dephosphorylation.

In summary, these results indicate that phosphorylation does not substantially alter either the predicted 9-mer binding core or the binding likelihood, reinforcing our previous observation that phosphosite localization preferences coincide with those of unmodified S/T/Y residues. While dephosphorylation may moderately weaken phosphopeptide–HLA-II interactions in some cases, according to the developed model, the overall impact is limited.

### A comment on phosphosite localization certainty

Our analyses thus far have not accounted for the ambiguity in phosphosite localization, as indicated by the AScore (or Ambiguity Score) parameter computed by PEAKS. A more detailed examination of this aspect is provided in the **Supplemental Note S1**. Briefly, in this examination, we observed that model performance does not improve when exclusively trained on high-confidence sites data (ligands with AScore > 13), suggesting limited sensitivity to localization certainty as annotated by PEAKS. Additionally, attempts to increase the accuracy of phosphosite localization annotation either by using phosphosite prediction tools, such as NetPhosPan^43^ or PhosphoLingo^44^, or by overlapping with public phosphoproteomics databases demonstrated, in all cases, a very limited recovery (i.e. only a small fraction of the observed phosphosites in the identified phospholigands were either predicted to be phosphorylated or matched to entries in the phosphoproteomics databases). In the case of phosphosite predictions, sequence composition analysis revealed biases in phosphosite prediction tools, which tend to favor cytosolic/nuclear kinase motifs. Therefore, these tools are of limited utility for our purposes, as most of the identified HLA-II phospholigands are derived from the endolysosomal pathway (as predicted by DeepLoc-2.0^45^). Together, these findings reveal that current phosphorylation prediction tools and databases are restricted by biases and specific dataset characteristics, emphasizing the need for tailored approaches to achieve accurate phosphosite localization assignment.

### NetMHCIIphosPan predictions on phosphopeptides identified by FragPipe

To further evaluate the generalizability of NetMHCIIphosPan and investigate the limited overlap between phospholigands identified by PEAKS and MaxQuant, we re-analyzed the Racle and Racle_DR datasets using FragPipe (v23.1, MSFragger v4.3, Philosopher v5.1.2^18,28,29^; for more details refer to Methods). These datasets were selected for analysis as they yielded the highest number of phosphopeptide identifications among all samples, as previously shown in **Fig. 1**. We further reran the PEAKS analyses with adjusted search parameters, including a 1% FDR and a reduced number of variable PTMs, to allow for a better comparison between the two analyses (for details refer to Methods). Comparing the sets of unmodified peptides identified by the two methods demonstrates a very high concordance between the two approaches both in terms of peptide yield (242,247 and 261,781 unique peptides identified by PEAKS and FragPipe, respectively) and overlap of 86% (with respect to the minority set). (see **Supplemental Fig. S11A**). However, when focusing on the phosphopeptides, the concordance between the two methods is greatly reduced. Here, the overlap is of 29% (with respect to the minority set), and the overall peptide yield is substantially lower when considering FragPipe compared to PEAKS, with 3320 and 5407 unique phosphorylated peptides identified by FragPipe and PEAKS, respectively, see **Supplemental Fig. S11B**. When phosphorylated residues are reverted to their unmodified forms, thereby disregarding both the number of phosphorylations and the exact phosphorylation sites when multiple S/T/Y acceptor sites are present within the same sequence, only a modest increase in overlap is observed (**Supplementary Fig S11C**).

To further assess the accuracy of the phospholigands identified by the two methods, we stratified the peptides into two subsets: peptides identified by both methods (“overlapping”), or peptides identified only by one of the two methods (“non-overlapping”), and next extracted prediction scores for the peptides using NetMHCIIphosPan. For peptides identified exclusively by FragPipe, predictions were generated as an external evaluation set, since these peptides were not included in model training. For peptides identified by PEAKS, which were used in model training, predictions were obtained through 5-fold cross-validation. As shown in **Supplemental Fig. S11D**, overlapping phosphopeptides displayed consistently high predicted presentation likelihoods, with close to 77% of the peptides scoring at %rank ≤ 1. Among non-overlapping peptides, both methods display a comparable performance with only 6-8% of the peptides being assigned a rank > 20%. In terms of rank values, PEAKS displays a small advantage over FragPipe, with 75% of the peptides being predicted with a rank ≤ 5%. In comparison, this value is 67% for FragPipe. Returning to the peptides identified using PEAKS with the search criteria applied to extract the training data for NetMHCIIphosPan (among other things relaxed to an FDR of 5% and an increased number of PTMs (refer to Methods and **Fig. 1**)), we show in **Supplemental Fig. S11E**, the distribution of percentile rank values for the two peptide subsets; the subset identified using the search criteria with the FDR of 1%, and the subset only identified using the relaxed settings. The results of this demonstrate a very similar %rank distribution of the two subsets, with less than 5% of the peptides in both datasets being assigned a rank > 20%, and more than 80% of the peptides being assigned a rank value less than 5%.

Together these results suggest that despite a rather low concordance between the identified sets of phosphorylated peptides between the FragPipe and PEAKS methods, the HLA binding patterns contained within the different datasets is consistent, allowing a method trained on the PEAKS data to accurately predict peptides identified by FragPipe. Further, the similarities between the percentile rank values for the peptides identified using an FDR of 1% or 5%, supports that the impact of the increased false discovery imposed by the relaxed FDR rate is very minor if at all present.

### Benchmarking against other earlier methods: MixMHC2pred-1.3

As a final test, the NetMHCIIphosPan method was benchmarked against MixMHC2pred-1.3^24^ employing an independent immunopeptidome dataset of 2058 phospholigands obtained from Esophageal adenocarcinoma (EAC) samples (11 cell lines) and Monocyte-derived dendritic cells pulsed with a recombinant SARS-CoV-2 Spike protein vaccine candidate (5 cell lines). Note, that the overall peptide yield from the EAC samples was poor given their low HLA-II expression (for details on this dataset refer to Materials and Methods). As explained before, we evaluated the performance with the ROC-AUC, ROC-AUC 0.1, PPV and PR-AUC metrics. Note that the external datasets only cover DRB1 and a few DRB3/4/5 alleles. Also, it is important to stress that, in contrast to NetMHCIIphosPan, MixMHC2pred-1.3 can only make predictions for alleles present in its training dataset. To accommodate for this limitation, the benchmark was limited to consider only the HLAs available in MixMHC2pred-1.3. Of note, when evaluating performance using this external benchmark, we ensured no overlap between the training dataset of our model (or NetMHCIIpan-4.3) and the independent evaluation data (for more details refer to Materials and Methods). In contrast, MixMHC2pred-1.3 was evaluated without controlling for such overlap, potentially resulting in inflated performance estimates due to data leakage.

**Fig. 8** presents the benchmark results, showing that NetMHCIIphosPan outperformed MixMHC2pred-1.3 across all metrics for the two external datasets (all P<0.0001; for performance evaluation details refer to Methods section). Remarkably, performance values for the EAC dataset remain low, particularly for ROC-AUC 0.1, PR-AUC and PPV, compared to the cross-validation results (**Figs. 2** and **4**), suggesting a lower quality for this dataset compared to the training datasets. **Supplemental Fig. S12** extends the comparison shown in **Fig. 8** by including prediction results from NetMHCIIpan-4.3, evaluated on three “phospho-equivalent” datasets. Additionally, **Supplemental Fig. S13** reports prediction performance across the full allele set, excluding MixMHC2pred-1.3, which supports only a limited set of alleles, as previously noted.

**Fig. 8.**
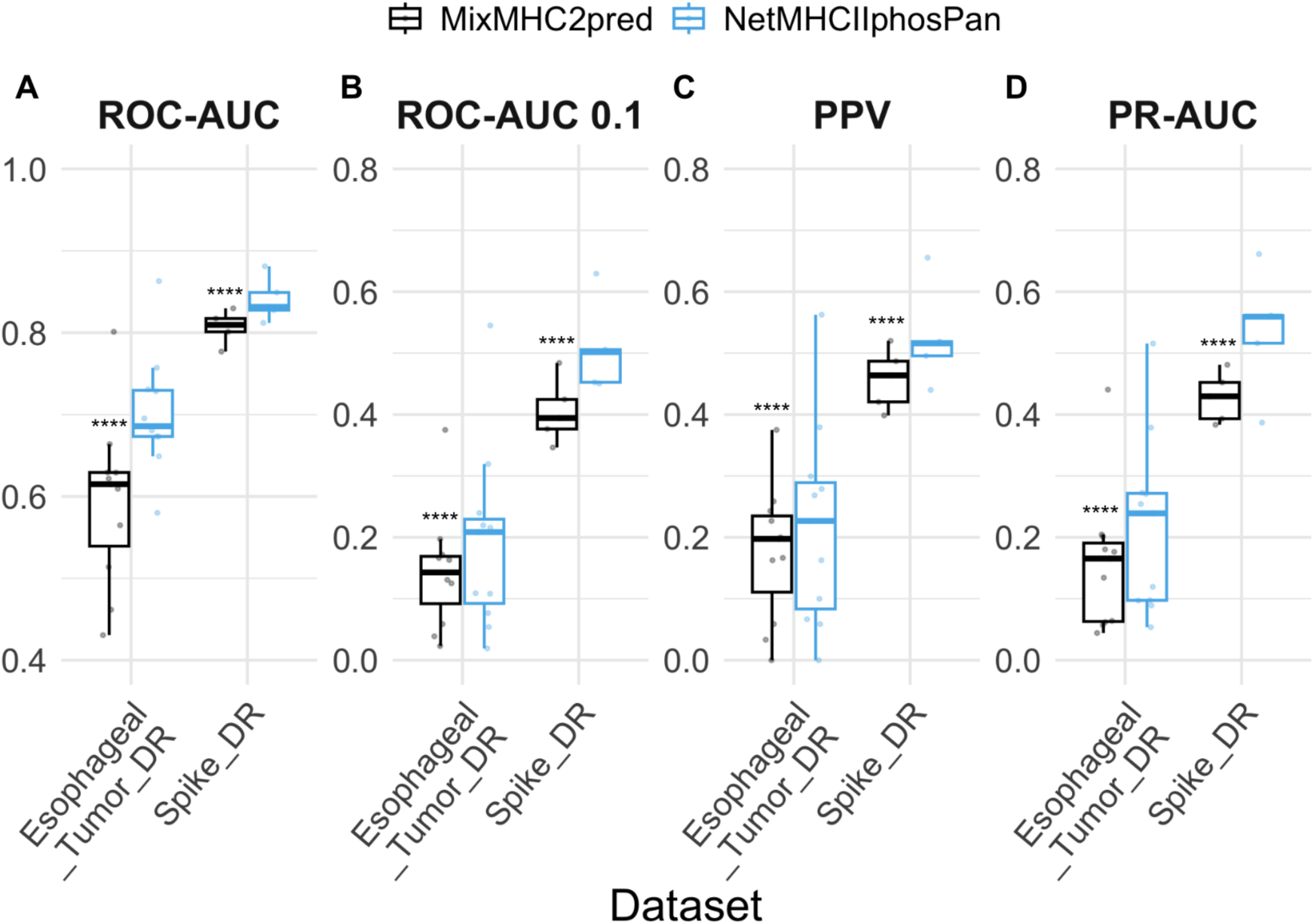
Performance of NetMHCIIphosPan and MixMHC2pred-1.3 methods on two phospholigand external evaluation datasets covering HLA-DR alleles. Performance metrics included are (**A**) ROC-AUC, (**B**) ROC-AUC0.1, (**C**) PPV and (**D**) PR-AUC. ROC-AUC refers to the area under the Receiver Operating Characteristic curve; ROC-AUC 0.1 is the partial AUC integrated up to a FPR of 10%; PR-AUC is the area under the Precision-Recall curve; and PPV (Positive Predictive Value) is calculated as described in Materials and Methods. Performance is evaluated in a per-cell line manner from the concatenated test sets. Metrics were computed for cell lines with at least 10 phospholigands (N=5 cell lines for “Spike_DR” and N=11 cell lines for “Esophageal_Tumor_DR”). The center line inside the box indicates the median value of the plotted metric and the box covers the interquartile range. The whiskers extend to, at most, 1.5-fold of the interquartile range. The individual data points in the jitter plots correspond to different cell lines. A one-tailed binomial test was employed for model performance comparison, and p-values were estimated using a bootstrap analysis, for more details refer to Materials and Methods. (**** P<0.0001). Y-axis ranges differ between panels **A** and **B**-**D**.

The NetMHCIIphosPan method is made available as a web server and stand-alone tool at https://services.healthtech.dtu.dk/services/NetMHCIIphosPan-1.0. The tool has functionality like the other NetMHC-tools and allows for the submission of either phosphopeptides or proteins with annotated phosphosites, making predictions for any HLA class II molecule with a known sequence (for details refer to the web server instructions).

## Discussion

This study introduces NetMHCIIphosPan, a novel predictor for phosphorylated peptide binding to HLA class II molecules, developed through a tailored ML framework and an extensive reanalysis of MS-based immunopeptidomics data. By addressing critical challenges such as data quality, and allele coverage in earlier phospholigand datasets, this work significantly advances the characterization of HLA class II phospholigand binding motifs and predictive modeling.

An important aspect of this study is the shift from a model-centric ML approach to a data-centric ML approach. In model-centric ML, the focus is on continuously improving the model itself while keeping the training data fixed. In contrast, data-centric ML focuses on improving the quality of the training data while holding the model architecture constant^46^. The initial approach to improve HLA-II phospholigand prediction followed a model-centric strategy, where the NNAlign_MA algorithm was trained using MaxQuant-identified datasets (filtered using prior knowledge of HLA-II unmodified peptide binding). However, this approach yielded suboptimal results due to the deficiencies of the training data described here. This limitation prompted a shift to a data-centric approach, prioritizing improvements in data quality by reprocessing raw MS datasets with a tailored PEAKS workflow. A key observation from this data-centric shift was a superior performance for identifying phosphorylated HLA class II ligands in terms of both peptide quantity and quality. Across datasets, the new workflow recovered 33% to 246% more phosphopeptides.

Using these re-annotated datasets, we developed predictive models using the NNAlign_MA framework. Here, the superior quality of the new data was manifested not only in terms of improved predictive performance metrics, but also in terms of a substantial reduction in the number of peptides classified as HLA-irrelevant contaminants. Investigating the identified HLA binding motifs, the model trained on the novel data further demonstrated superior consistency compared to models built with MaxQuant-identified phosphopeptides. Here, the median Pearson Correlation Coefficient (PCC) between motifs for phosphorylated and unmodified ligands was 0.39 for the M1 and M2 models, compared to 0.82 for the M3 model, highlighting the superior ability of PEAKS-based models to capture binding preferences. Additionally, incorporating unmodified ligands into the training dataset and allowing peptides to bind in the inverted mode further refined the predictive model, which achieved superior performance values across all metrics (ROC-AUC, ROC-AUC0.1, PPV and PR-AUC) compared to the baseline model. As expected, motif consistency was increased for HLA-DP alleles, from a median PCC of 0.86 to 0.91. For HLA-DQ alleles, the median PCC improved from 0.60 to 0.77, suggesting that the inclusion of unmodified ligands helped to mitigate the limited coverage of the HLA-DQ phospholigand data.

Phosphosite localization analysis revealed nuanced enrichment and depletion patterns across HLA class II molecules. Regarding phosphosite placement relative to the predicted binding core, approximately 20% of phosphosites are found in both the N- and C-terminal flanking regions, contradicting the findings of Solleder *et al.* who reported a higher occurrence of phosphosites in the C-terminus.

Within the binding core, we observed clear differences from HLA class I phospholigands, which tend to favor phosphoresidues at P4, particularly for pS^41,47^. In this regard, Solleder *et al.* made two key observations: (1) a predominance of phosphosites at P5 within the binding core, and (2) a low frequency of phosphoresidues at the anchor position P1. Our findings diverge from the first observation, as we did not detect an enrichment of phosphorylated residues at P5. Instead, our analysis did reveal a marked depletion of pS and pT at P1 across most DRB1, DRB3/4/5, and HLA-DP alleles, along with a general tendency for phosphosites to be disfavored at P1 for most alleles and phosphorylation types. However, an exception to this trend is pY and DRB1 alleles, where a small proportion showed an enrichment at P1. Finally, consistent with Solleder *et al.* we found that the binding motifs of unmodified peptides closely resemble those of phosphopeptides. Our current model predicts that phosphosite location within the binding core is strongly correlated with binding motif preferences for unmodified S/T/Y residues, reinforcing the notion that HLA class II phospholigand motifs are primarily shaped by the inherent binding preferences of unmodified ligands.

As mentioned earlier, a key challenge for annotating phosphopeptides is the identification of the phosphorylation site. To address this, we investigated if phosphosite recovery and thus, localization certainty, could be improved by using prediction tools such as NetPhosPan and PhosphoLingo, or by cross-referencing with public phosphoproteomics datasets. However, our results revealed the limited potential of such approaches, as only a small fraction of the phosphosites identified in our MS spectra overlapped with high-scoring predicted sites or reported entries. We suggest that this discrepancy arises from dataset-specific characteristics or biases. HLA-II phospholigands predominantly originate from endolysosomal pathway proteins, whereas most phosphosite prediction tools and databases are primarily based on proteins from the cytosol or nucleus. As a result, these tools have limited applicability in our context, highlighting the need for refined approaches to achieve accurate phosphosite localization.

To further assess phosphopeptide identification quality and investigate the limited overlap between the peptides identified using the PEAKS workflow compared to MaxQuant, we re-analyzed the Racle datasets using FragPipe (v23.1). The FragPipe analysis yielded an overall substantial overlap with PEAKS. This overlap was however considerably reduced for phosphorylated peptide identifications compared to the overlap found for unmodified peptides. The vast majority of overlapping phosphopeptides were consistently predicted as strong HLA-II binders by NetMHCIIphosPan. Notably, among non-overlapping identifications between the two methods, the FragPipe-unique phosphopeptides displayed an HLA binding profile very similar to that of the PEAKS-unique phosphopeptides both containing more than 75% of the peptides with a NetMHCIIphosPan predicted percentile rank score of 5 or less. These results support the high confidence of FragPipe and PEAKS outputs and highlight their reliability in recovering biologically relevant phospholigands. However, the fact that the overlap in the identified phosphopeptide space between the two methods is much reduced compared to that observed for unmodified peptides, suggests that neither method is performing optimally when applied in the context of PTM identification, and that further refinement of the tools is needed. Of note, for the workflow developed here, we have opted for using a relaxed FDR threshold of 5% for the peptide identification. This naturally comes with a risk of false positive ligand identifications. However, when comparing the peptides identified using an FDR of 1% or 5%, we find very similar predicted binding score distributions, suggesting that, at least when it comes to learning the patterns associated with HLA antigen presentation, the relaxed false discovery rate has very minor impact, if any at all. It is however essential to underline that the application of such a relaxed FDR might not be recommendable in cases where the workflow is applied for direct antigen discovery.

Finally, when benchmarking NetMHCIIphosPan against existing tools, including NetMHCIIpan-4.3 and MixMHC2pred-1.3, it demonstrated superior predictive accuracy. Although external benchmarks were limited to DRB1 and some DRB3/4/5 molecules, the results highlight the robustness of NetMHCIIphosPan and suggest opportunities for further validation on expanded datasets covering HLA-DP and HLA-DQ alleles.

By integrating data-centric ML principles with comprehensive MS datasets, NetMHCIIphosPan offers a state-of-the-art method for analyzing phosphorylated peptide binding to HLA class II molecules. This work underscores the importance of using high-quality datasets in training predictive modeling and provides new insights into the presentation of phosphorylated peptides, setting the stage for future studies exploring their role in immune recognition, autoimmune diseases, and cancer. Further, we envision that similar approaches combining refined MS peptide identification workflows with tailored machine learning methods could be applied to other PTMs, to cast light on their roles in HLA antigen presentation.

## Abbreviations

HLA: Human Leukocyte Antigen
HLA-II: Human Leukocyte Antigen class II
MS: Mass Spectrometry
MHC: Major Histocompatibility Complex
ML: Machine Learning
PTM: Post-Translational Modification
SA: Single-Allele
MA: Multi-Allele
BA: Binding Affinity
EL: Eluted Ligand
PH: Phosphopeptide
UM: Unmodified peptide
FPR: False Positive Rate
ROC-AUC: Receiver Operating Characteristic - Area Under the Curve
ROC-AUC 0.1: Receiver Operating Characteristic - Area Under the Curve integrated up to a 0.1
FPR PPV: Positive Predictive Value
PR-AUC: Precision-Recall - Area Under the Curve

## Data Availability Statement

The MS proteomics data, generated in this study for the external benchmark, have been deposited in the ProteomeXchange Consortium via the PRIDE^48^ partner repository with the dataset identifier PXD060958. Furthermore, the training and evaluation data for NetMHCIIphosPan-1.0 is available on its web server at https://services.healthtech.dtu.dk/services/NetMHCIIphosPan-1.0, “Supplementary material” tab.

## Supporting Information

This article contains supporting information.

## Author Contributions

Conceptualization: M.N.

Methodology: M.N., S.K., N.T. and H.M.G.A.

Investigation (data analysis and model development): M.N. and H.M.G.A.

Investigation (re-analysis of raw MS datasets): S.K., H.Y., C.M.S. and N.T.

Investigation (wet-lab experiments): R.P., N.T., B.P. and A.S.

Visualization: H.M.G.A.

Supervision: M.N. and S.K.

Writing (original draft): H.M.G.A. and M.N.

Writing (review and editing): All authors

## Funding

Research reported in this publication was supported in part by the National Cancer Institute (NCI), under award number U24CA248138, the National Institute of Allergy and Infectious Diseases (NIAID), under award number 75N93019C00001, and the Cancer Research UK Centres Network Accelerator Award grant (C328/A21998).

## Ethics Statement

Most of the human data used in this study were obtained from the PRIDE database and/or previously published studies that had received appropriate ethical approval and documented informed consent from participants. In addition, the study that generated the “Esophageal tumor” dataset, included in the independent benchmark, was conducted in accordance with the Declaration of Helsinki and approved by the South Central–Oxford C Research Ethics Committee (REC reference: 16/SC/0015). The study protocol was registered under protocol number LUD2015-005, with EudraCT number 2015-005298-19 and IRAS project ID 194443. All subjects involved in this study provided written informed consent prior to participation.

## Competing Interests

The authors declare no competing financial interest.

## Acknowledgments

We thank the Southampton University Hospitals and Oxford University Hospitals for collecting the tumor samples and thank Prof. Xin Lu and Prof. Mark Middleton for providing resources and their support of the research project.

## Notes

### Competing Interest Statement

The authors have declared no competing interest.

