## Supporting_Information for "NetMHCIIphosPan: a machine learning tool for predicting HLA class II antigen presentation of phosphorylated peptides"

Heli M. Garcia Alvarez *et al.*

**This PDF file includes:**

Supplemental Figures S1 to S13
Legends for Supplemental Tables S1 to S5
Supplemental Note S1
References for Supplemental Note S1

**Other Supporing Information for this manuscript includes the following:**

Supplemental Tables S1 to S5

### Supplemental Figures

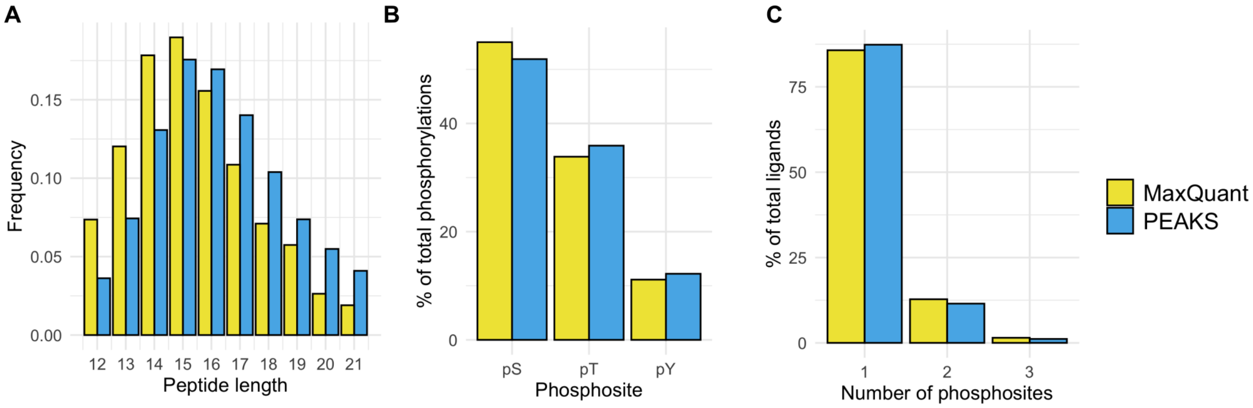


**Fig. S1**. **Overview of the phospholigand datasets identified by MaxQuant and PEAKS software.** (**A**) Peptide length distribution. (**B**) Phosphorylation type prevalence, considering all phosphosites found in modified peptides. (**C**) Distribution of the number of phosphosites per modified peptide. Abelin, Racle, PvanBalen, and Saghar (DR, DP, and DQ) cell lines are included within the PEAKS datasets.

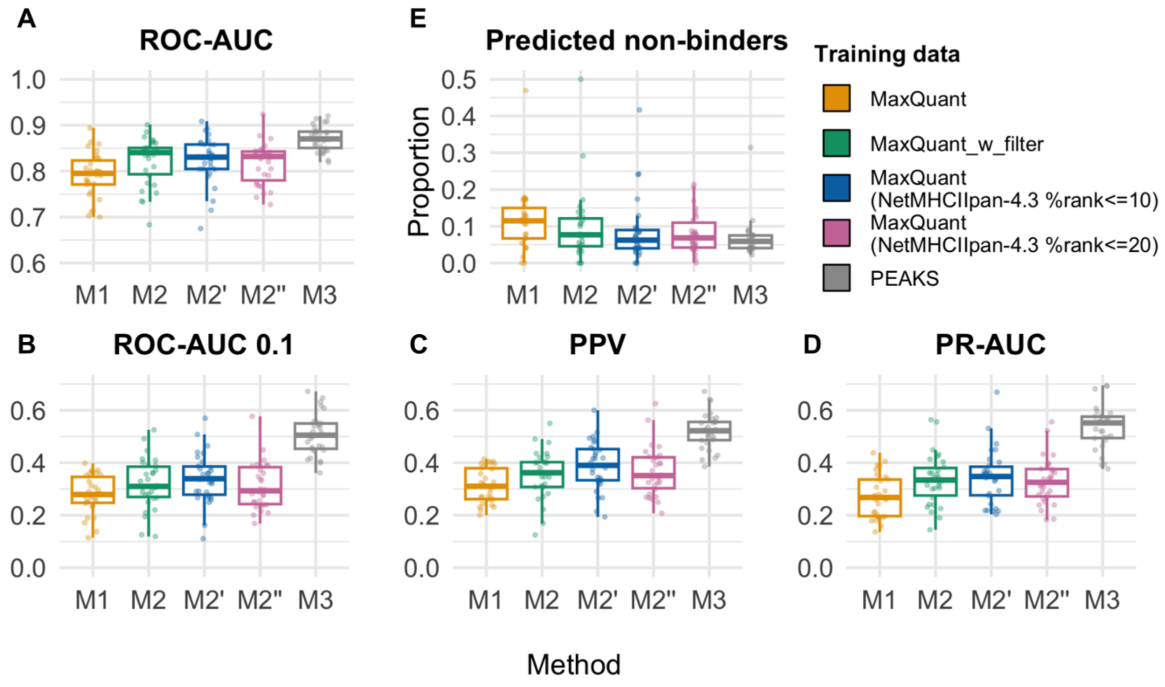


**Fig. S2. Five-fold cross-validation performance of the methods trained with phospholigands identified by MaxQuant and PEAKS. M1** refers to the model trained and evaluated on the complete MaxQuant dataset; **M2** refers to the model trained and evaluated on the MaxQuant dataset, pre-filtered according to MixMHC2pred-2.0 predictions (%rank≤10) (named "MaxQuant_w_filter"); **M2'** refers to the model trained and evaluated on the MaxQuant dataset, pre-filtered according to NetMHCIIpan-4.3 predictions (%rank≤10); **M2''** refers to the model trained and evaluated on the MaxQuant dataset, pre-filtered according to NetMHCIIpan-4.3 predictions (%rank≤20); **M3** refers to the model trained and evaluated on the complete PEAKS dataset. Performance metrics included are (**A**) ROC-AUC, (**B**) ROC-AUC0.1, (**C**) PPV and (**D**) PR-AUC. (**E**) Proportion of predicted non-binders (%rank>20) according to each of the methods. The center line inside the box indicates the median value of the plotted metric and the box covers the interquartile range. The whiskers extend to, at most, 1.5-fold of the interquartile range. The individual data points in the jitter plot correspond to different cell lines. The plots show the performance metrics calculated over Abelin and Racle datasets (N=30 cell lines), with 10 or more positive instances for all trained methods. Y-axis ranges differ between panels **A** and **B-D**.

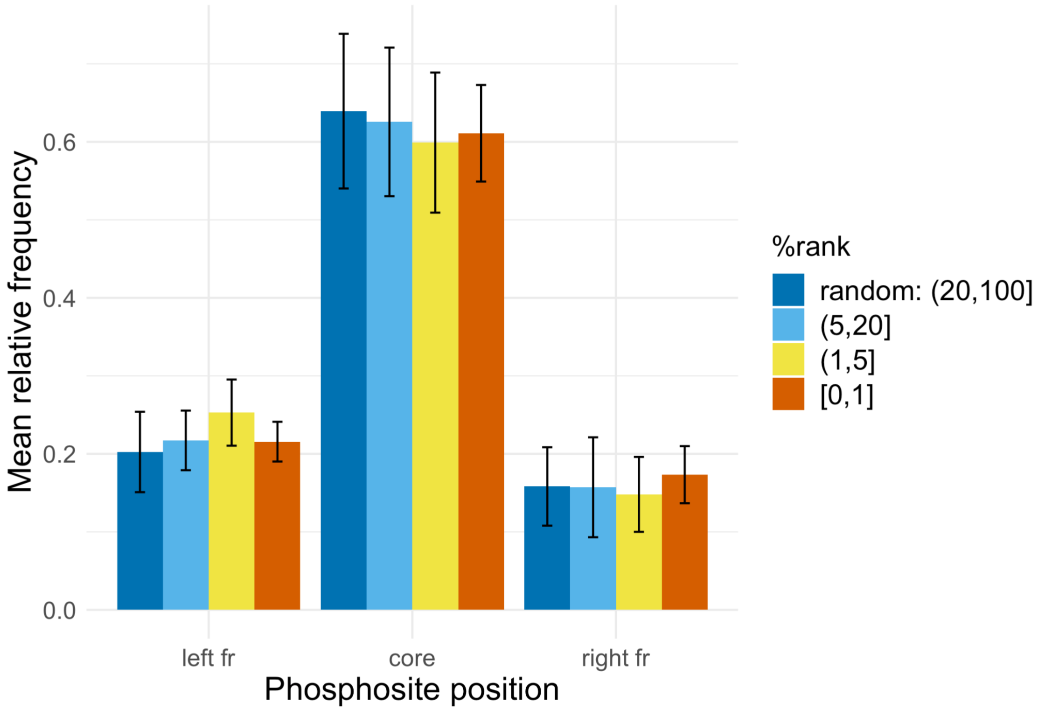


**Fig. S3**. **Phosphosite localization distribution for ligands and random peptides averaged across peptide lengths 13 to 17**. Yellow, orange and light blue bars correspond to ligands, while random peptides are depicted in blue. The error bars show the standard deviation. The predicted NetMHCIIphosPan binding core is 9-aa long, while the left and right flanking regions ("fr") correspond to fragments of a variable length outside the core.

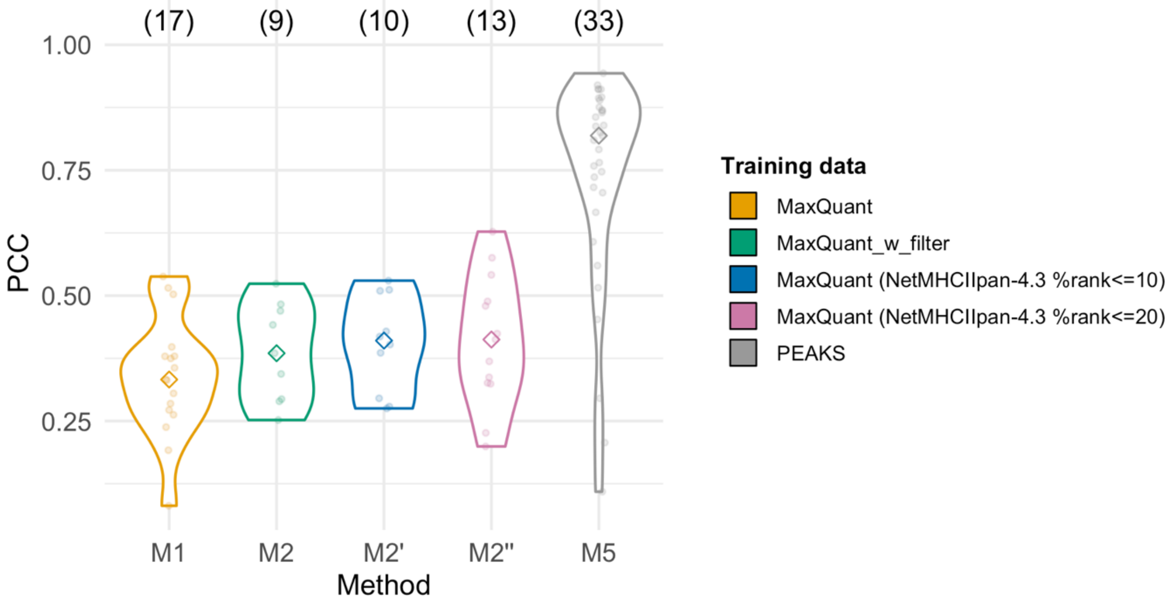


**Fig. S4**. **Concordance between unmodified and phospholigand binding motifs for HLA-II molecules with 30 or more peptides assigned during method training.** The methods shown here are the same as in **Fig S2**. PSFMs (position-specific frequency matrices) were computed to represent HLA-II binding motifs for unmodified and phospho-peptides, based on cross-validation predictions. In the case of phospho-peptides, phosphosites were ignored and frequencies per position were renormalized to sum up to 1. For unmodified peptides, predictions from NetMHCIIpan-4.3 were employed. To compare binding motifs for a given allele, PCCs (Pearson Correlation Coefficient) were calculated between flattened unmodified and phospho-peptide PSFMs. The violin plots show the distribution of PCCs for each of the trained methods (each data point corresponds to a different HLA-II molecule, the diamond indicates the median PCC, and above each plot, the analyzed number of HLA-II molecules is noted).

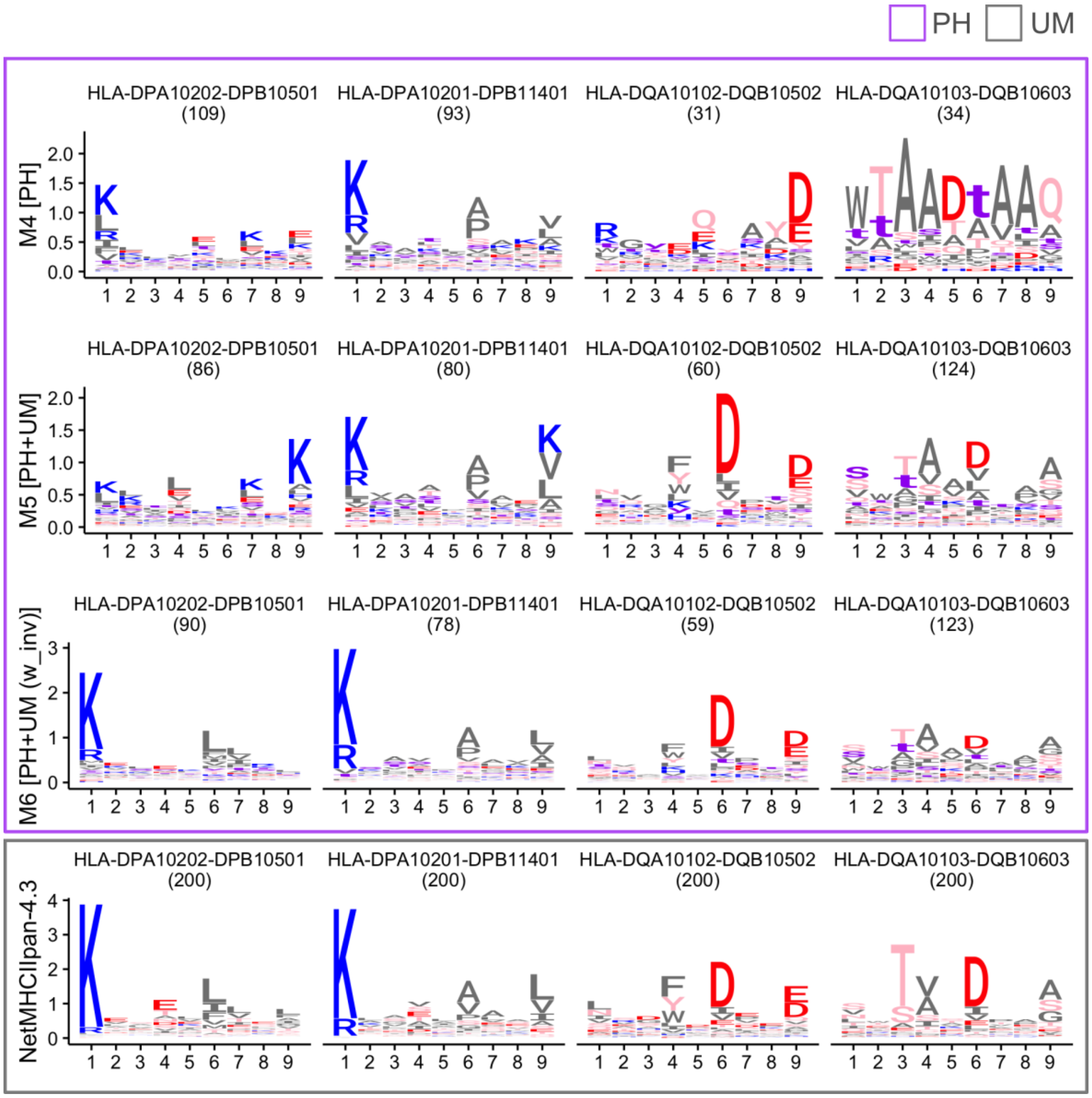


**Fig. S5**. **Sequence logos for two selected HLA-DP and HLA-DQ molecules**. Logos were generated from cross-validation predictions, following the analysis performed in **Fig. 4(E)**. The first three rows contain logos illustrating phospholigand ("PH", outlined in violet) binding motifs for each of the trained methods, M4 to M6. The fourth row shows logos representing unmodified ("UM", outlined in grey) HLA-II ligand binding motifs according to NetMHCIIpan-4.3 cross-validation predictions (200 peptides were randomly sampled per HLA-II molecule). Each logo displays in parenthesis the number of peptides employed for its generation. Amino acid color coding: basic in blue (RHK), acidic in red (DE), polar in pink (STNQYC), hydrophobic in dark grey (GPLIVAMFW), and phosphosites in violet (sty).

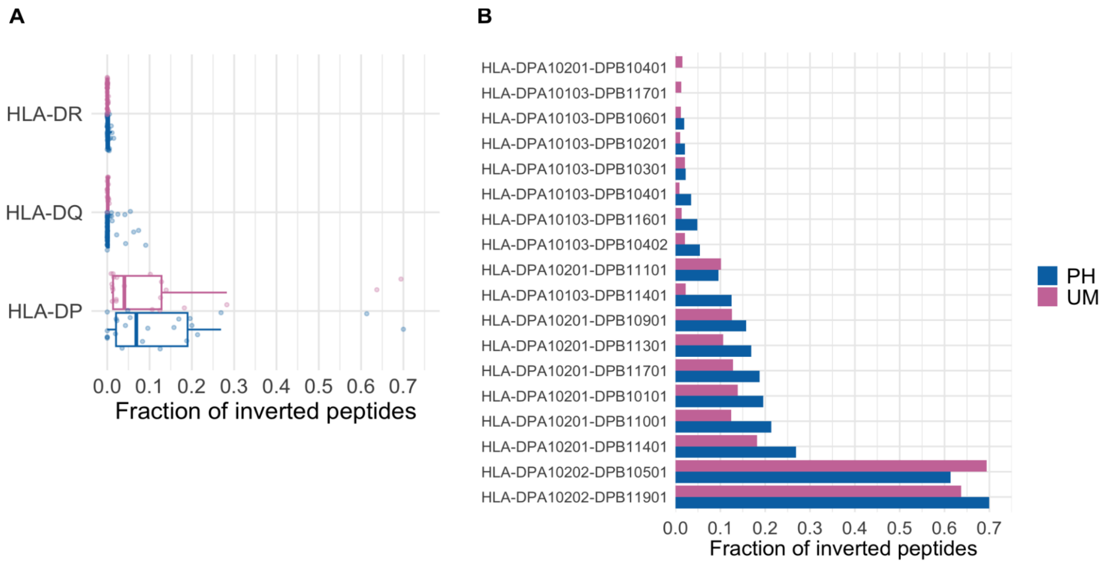


**Fig. S6. Fraction of peptides predicted to bind in the inverted mode according to the M6 method, trained on unmodified and phospholigands identified by PEAKS.** (**A**) Proportion of peptide inversion across HLA-II loci for predicted unmodified ligands ("UM") and phospholigands ("PH") from the method's training dataset. Peptides with a predicted percentile rank greater than 20% were discarded to focus the analysis on confident binders. The center line inside the box indicates the median and the box covers the interquartile range. The whiskers extend to, at most, 1.5-fold of the interquartile range. The individual data points in the jitter plots correspond to different HLA-II molecules. (**B**) Proportion of inverted peptides for HLA-DP molecules present both in the unmodified ligand and phospholigand datasets.

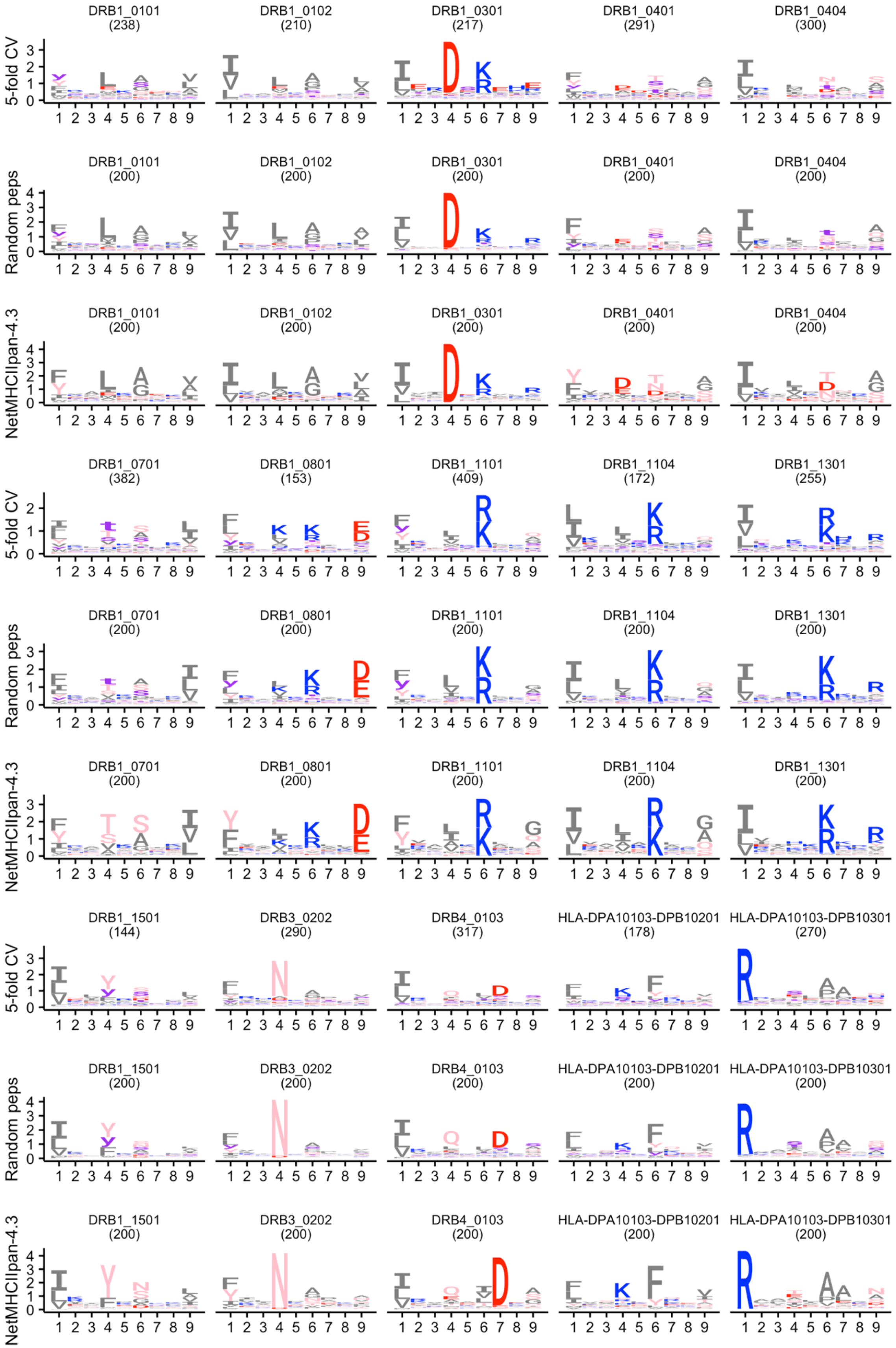
5-fold CV
Random peps
NetMHCIIpan-4.3
DRB1_0101 (238)
DRB1_0102 (210)
DRB1_0301 (217)
DRB1_0401 (291)
DRB1_0404 (300)
DRB1_0101 (200)
DRB1_0102 (200)
DRB1_0301 (200)
DRB1_0401 (200)
DRB1_0404 (200)
DRB1_0701 (382)
DRB1_0801 (153)
DRB1_1101 (409)
DRB1_1104 (172)
DRB1_1301 (255)
DRB1_0701 (200)
DRB1_0801 (200)
DRB1_1101 (200)
DRB1_1104 (200)
DRB1_1301 (200)
DRB1_1501 (144)
DRB3_0202 (290)
DRB4_0103 (317)
HLA-DPA10103-DPB10201 (178)
HLA-DPA10103-DPB10301 (270)
DRB1_1501 (200)
DRB3_0202 (200)
DRB4_0103 (200)
HLA-DPA10103-DPB10201 (200)
HLA-DPA10103-DPB10301 (200)

**(Figure continued)**

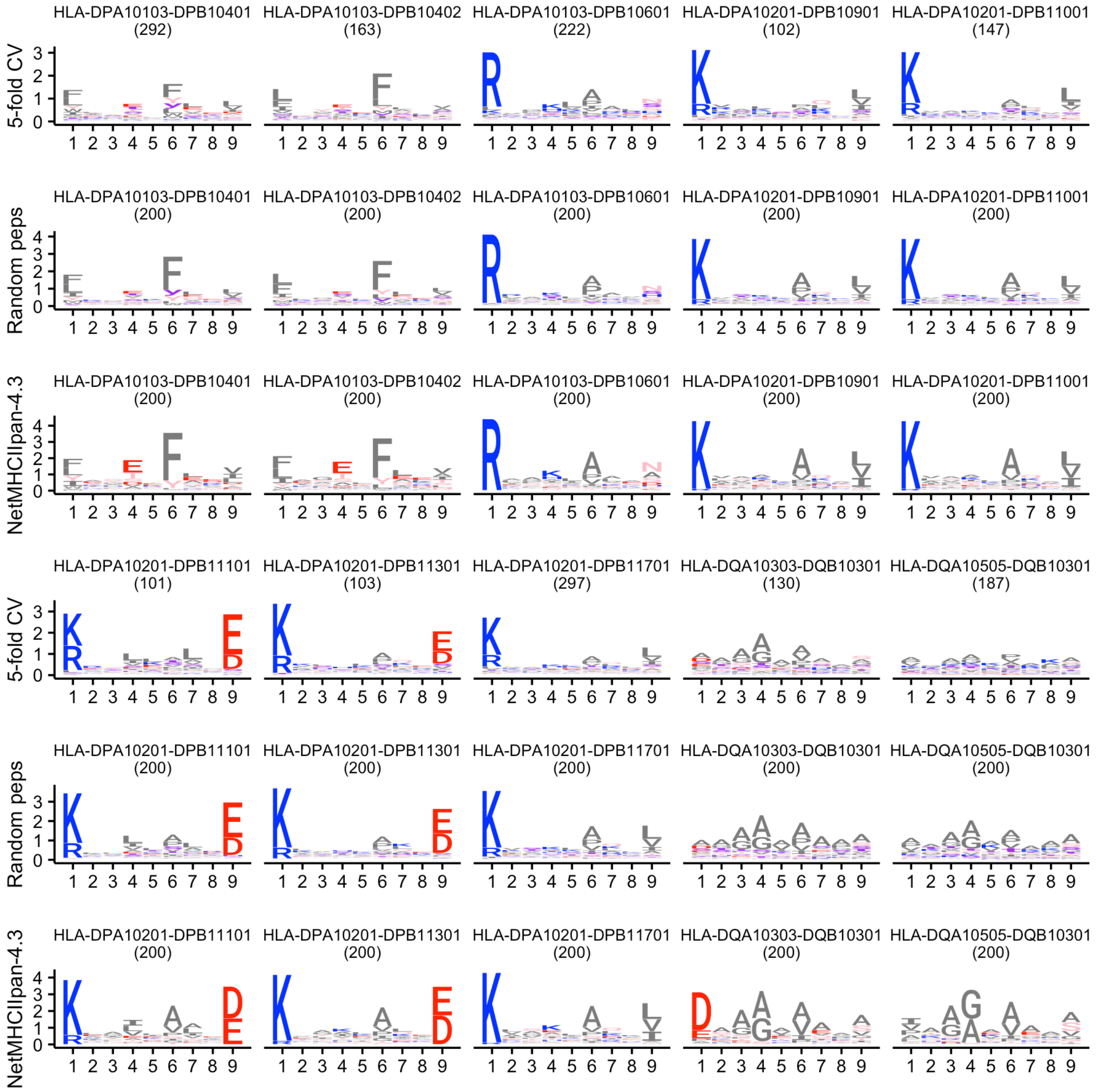


**Fig. S7. Sequence logo representations of HLA-II binding preferences for unmodified and phosphorylated peptides.** The plot includes the top 25 HLA-II alleles with the highest number of phospholigands assigned during NetMHCIIphosPan training. The first two rows correspond to NetMHCIIphosPan predictions from 5-fold cross-validation ("5-fold CV") and the top 0.1% scoring predictions from a set of 200,000 random natural phosphopeptides ("Random peps"). For comparison, logos for NetMHCIIpan-4.3 (third row) were generated using the top 0.1% scoring predictions from a set of 200,000 random natural (unmodified) peptides. In all cases, phosphopeptides with predicted %rank > 5 or lacking a phosphosite within the predicted binding core were excluded. Amino acid color coding: basic in blue (RHK), acidic in red (DE), polar in pink (STNQYC), hydrophobic in dark grey (GPLIVAMFW), and phosphosites in violet (sty). The number of peptides used to generate each logo is indicated in parentheses.

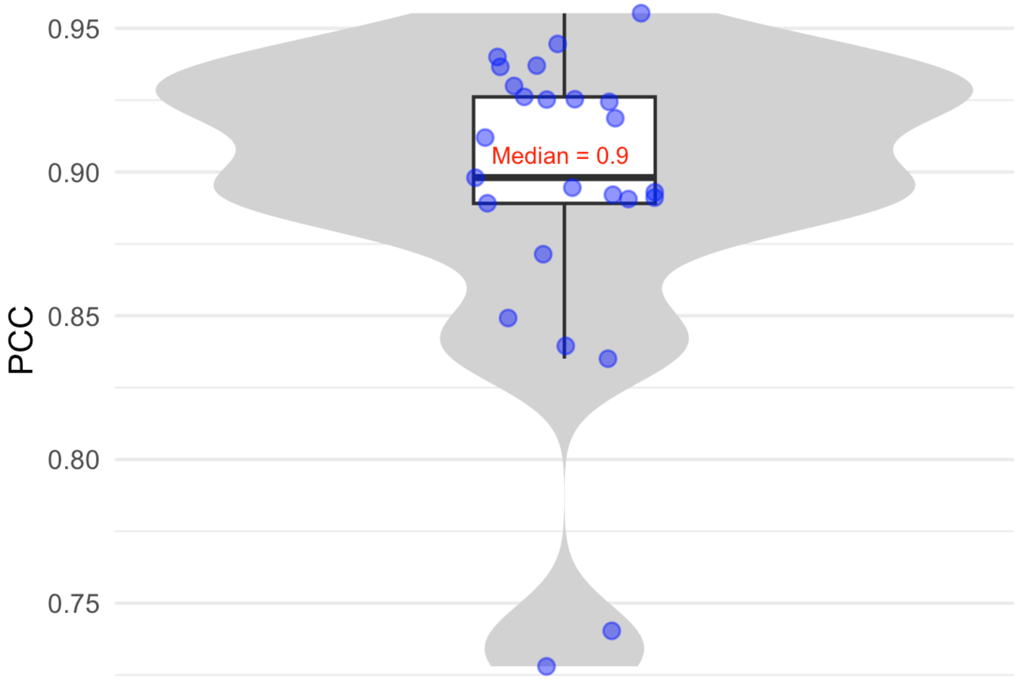


**Fig. S8**. **Distribution of PCCs computed between the flattened position-specific frequency matrices (PSFMs) corresponding to NetMHCIIphosPan logos constructed on random phosphopeptides and five-fold cross-validation phospholigands (Fig. S7)**. The center line inside the box indicates the median PCC and the box covers the interquartile range. The whiskers extend to, at most, 1.5-fold of the interquartile range. The individual data points in the jitter plots correspond to different HLA-II molecules (N=25).

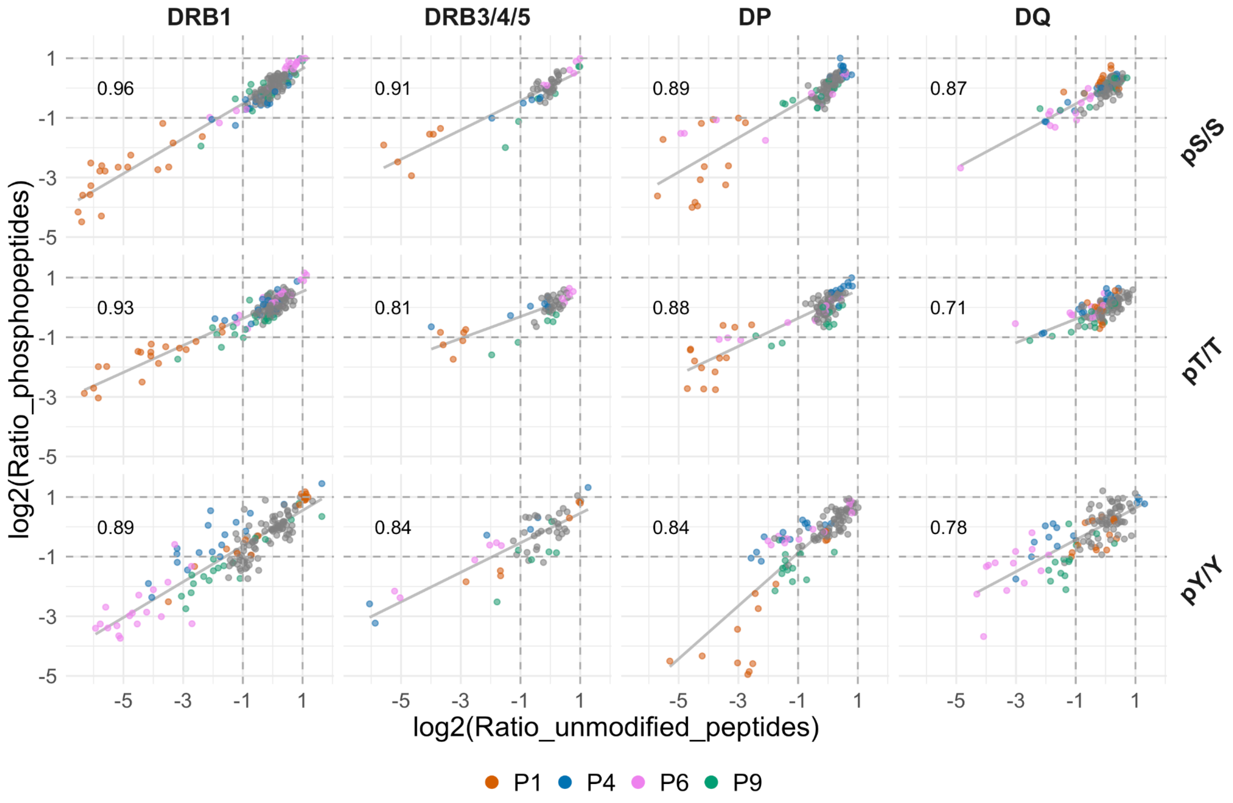


**Fig. S9. Correlation between the log2 ratio for random natural phosphopeptides and unmodified peptides, discriminated by HLA-II loci and amino acid (pS/S, pT/T, and pY/Y).** The ratio is calculated between the relative frequency of binders vs. non-binders for all 9 binding core positions, as in **Fig. 7**. Each data point corresponds to a given allele and position (P1 to P9). Data points corresponding to typical anchor positions for unmodified peptides, P1, P4, P6, and P9 are colored in orange, blue, pink, and green respectively. All other positions are shown in grey. The upper left corner of each subplot displays the Pearson Correlation Coefficient between the two plotted variables.

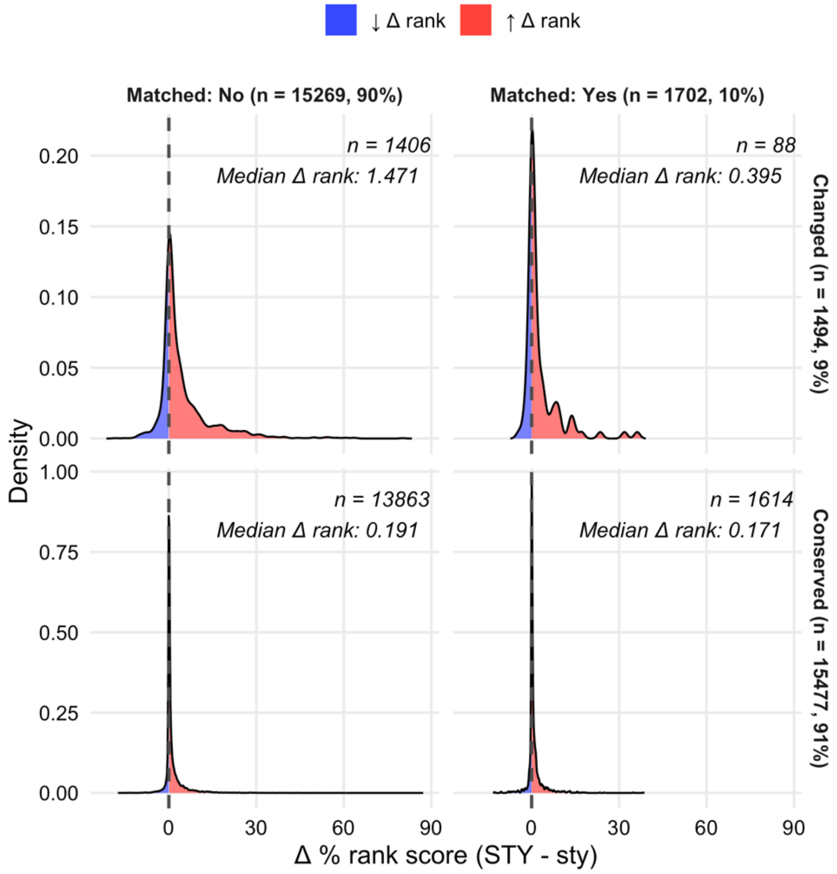


**Fig. S10. Impact of dephosphorylation on NetMHCIIphosPan predicted percentile rank scores for HLA-II phospholigands.** This figure shows the distribution of changes in predicted percentile rank scores (Δ %rank = STY − sty), comparing unmodified (STY) and phosphorylated (sty) versions of peptides. All phospholigands analyzed here were included in the final training of the NetMHCIIphosPan method, except for those considered likely contaminants (%rank sty > 20), which were excluded from the analysis (10.8%). For each, a dephosphorylated version was generated by reverting the phosphorylated residue (pS/pT/pY) to its unmodified form (S/T/Y), and both forms were evaluated using cross-validation. Peptides are grouped by whether their unmodified version is present in the unmodified eluted ligand training data (Matched) or not (Not Matched), and whether the predicted 9-mer binding core remained the same (Conserved) or changed (Changed) between the phosphorylated and unmodified forms. Red density regions (↑ Δ rank) indicate reduced predicted binding after dephosphorylation, while blue regions (↓ Δ rank) indicate improved binding. Median Δ rank values and peptide counts are reported for each group.

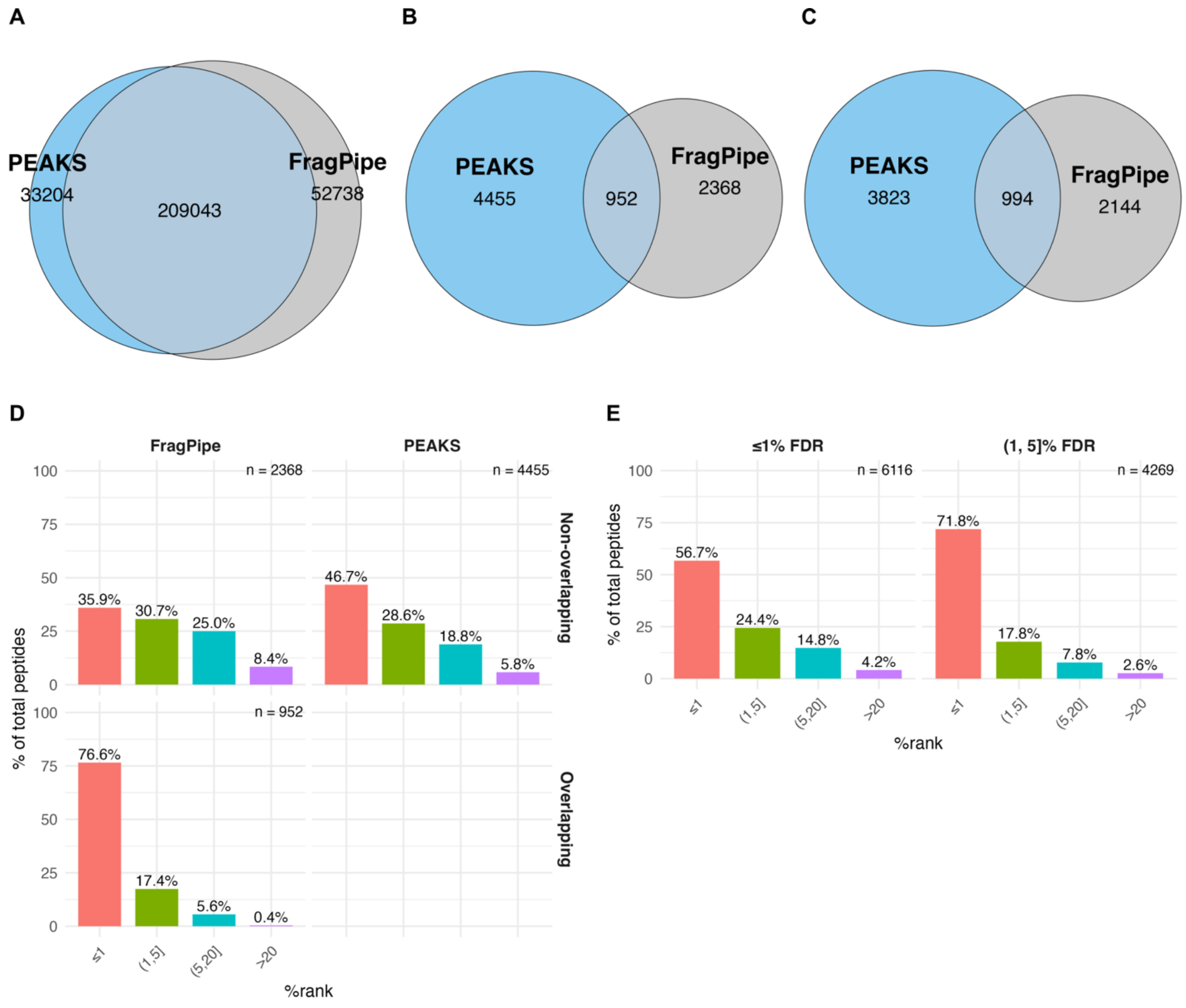


**Fig. S11. Overlap and NetMHCIIphosPan predicted HLA-II presentation of phosphopeptides identified by PEAKS and FragPipe in the Racle datasets**. Venn diagram showing the overlap of (**A**) unique unmodified peptides, (**B**) unique phosphopeptides, (**C**) unique phosphopeptides reverting phosphorylated amino acids to their unmodified forms, identified by PEAKS and FragPipe for the Racle and Racle_DR datasets. (**D**) Predicted HLA class II presentation likelihoods for phosphopeptides identified by each tool. Phosphopeptides were grouped based on whether they were identified by both tools or exclusively by one. For peptides with a 9-mer overlap with peptides placed in a given NetMHCIIphosPan training partition, predictions were made using the neural network ensemble that was not trained on that specific partition. If no overlap was found, a random neural network ensemble was selected for predictions. (**E**) Predicted HLA class II presentation likelihoods for training dataset PEAKS identified phosphopeptides grouped by FDR. For both (**D**) and (**E**), predicted %rank scores indicate the likelihood of HLA-II presentation, with lower values corresponding to stronger predicted binders. The number of peptides in each category is indicated in the top right corner of each panel.

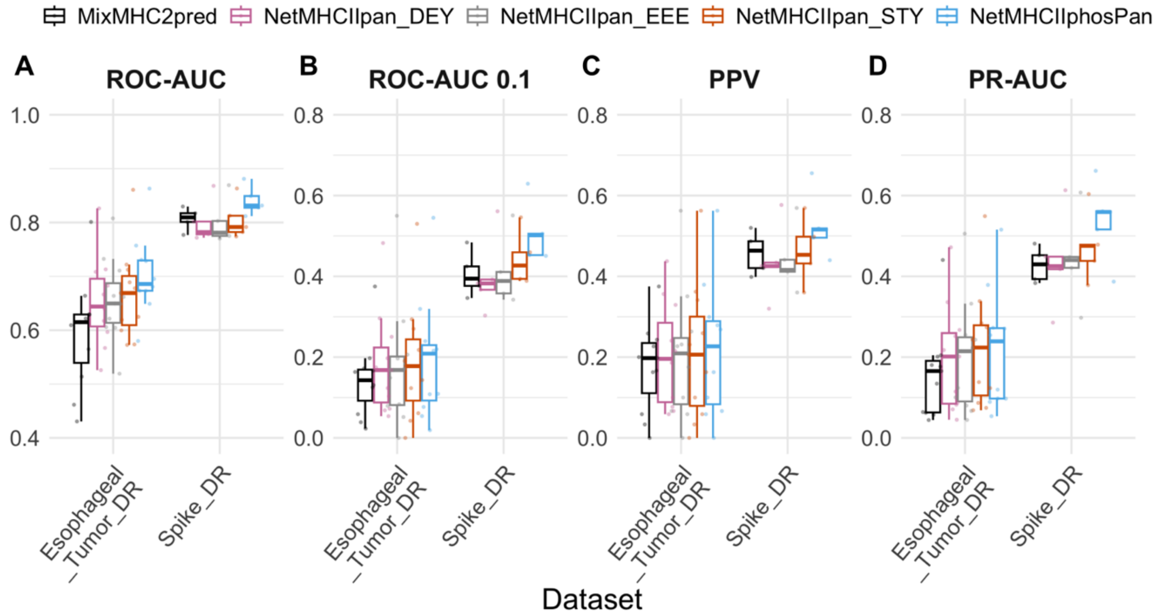


**Fig. S12**. **Performance of NetMHCIIphosPan, NetMHCIIpan-4.3, and MixMHC2pred-1.3 methods on two external evaluation datasets.** Predictions covered the reduced subset of HLA-DR alleles supported by MixMHC2pred-1.3 (N=5 cell lines for "Spike_DR" and N=11 cell lines for "Esophageal_Tumor_DR"). Performance metrics included are (**A**) ROC-AUC, (**B**) ROC-AUC0.1, (**C**) PPV and (**D**) PR-AUC. Performance is evaluated in a per-dataset manner from the concatenated test sets (for details see Materials and Methods). The center line inside the box indicates the median value of the plotted metric and the box covers the interquartile range. The whiskers extend to, at most, 1.5-fold of the interquartile range. The individual data points in the jitter plots correspond to different cell lines. Y-axis ranges differ between panels **A** and **B-D**.

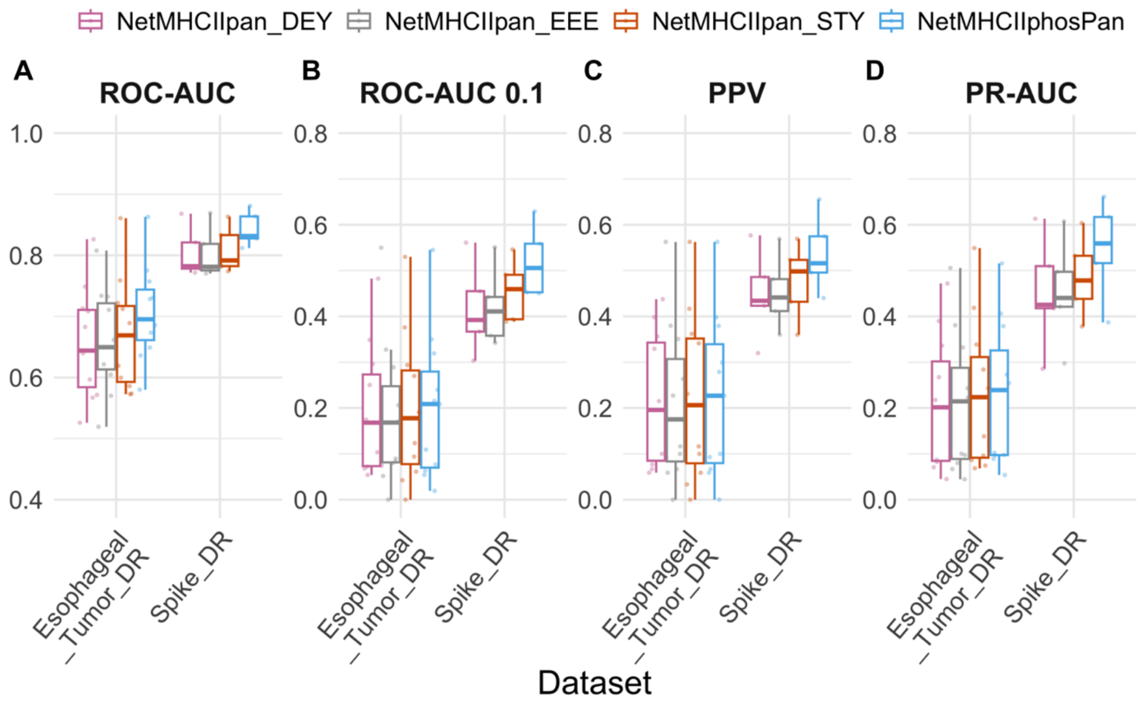


**Fig. S13**. **Performance of NetMHCIIphosPan and NetMHCIIpan-4.3 on two external evaluation datasets.** Predictions covered the full list HLA-DR alleles (N=5 cell lines for "Spike_DR" and N=11 cell lines for "Esophageal_Tumor_DR"). Performance metrics included are (**A**) ROC-AUC, (**B**) ROC-AUC0.1, (**C**) PPV and (**D**) PR-AUC. Performance is evaluated in a per-dataset manner from the concatenated test sets (for details see Materials and Methods). The center line inside the box indicates the median value of the plotted metric and the box covers the interquartile range. The whiskers extend to, at most, 1.5-fold of the interquartile range. The individual data points in the jitter plots correspond to different cell lines. Y-axis ranges differ between panels **A** and **B-D**.

### Legends for Supplemental Tables

**Table S1: List of identified phosphopeptides from the training datasets (Abelin, Racle, Saghar DR/DP/DQ, and PvanBalen_DP) generated using the PEAKS software with the initial set of search parameters** (including a 5% FDR). In addition, phosphopeptides identified from the Racle datasets using both FragPipe and PEAKS with the second set of search parameters (including a 1% FDR) are included for comparison. All listed phosphopeptides are 12–21 amino acid long.

**Table S2: List of identified phosphopeptides for the external benchmark datasets (Esophageal_Tumor_DR and Spike_DR) with the PEAKS software.**

**Table S3: Training dataset composition for each method presented in this study.** This table reports the number of positive instances (phospholigands), negative instances (random phosphopeptides), and the number of distinct cell lines included in the training dataset of each method described in this study.

**Table S4: Performance metrics computed for each of the trained methods in this study.** The table contains the per-dataset performance metrics (ROC-AUC, ROC-AUC 0.1, PPV and PR-AUC) reported for the methods shown in main Fig. 2, Fig. 4, Fig. 5, Fig. 8 and Supplemental Figs. S2, S12 and S13.

**Table S5: Phosphosite enrichment/depletion within the NetMHCIIphosPan predicted 9-mer binding core, computed for each binding core position, HLA-II molecule (N=50) and phosphosite type (pS/pT/pY), as shown in Fig. 7.** The table contains the calculated log2(Ratio) employed to construct Fig. 7.

### Tables are provided in separate files.

### Supplemental Note S1

A relevant point when considering the interpretation of the raw MS data is the definition of phosphorylation site localization. Peptides with multiple phosphate group acceptor sites, i.e. more than one serine, threonine, or tyrosine, suffer from a potential ambiguous phosphorylation site annotation. To address this, the PEAKS software computes for, each annotated phosphosite, a probabilistic metric called AScore, or ambiguity score, as defined by Beausoleil *et al.*[1]. This metric quantifies the probability of correct phosphosite localization based on the presence and intensity of site-determining ions in the MS2 spectra, as these ions vary between positional isomers. Considering this, we analyzed the NetMHCIIphosPan five-fold cross-validation percentile rank predictions for phospholigands and their associated AScores, as reported by PEAKS. For phosphopeptides with more than one phosphosite, the median AScore was calculated (see **Fig. 1**).

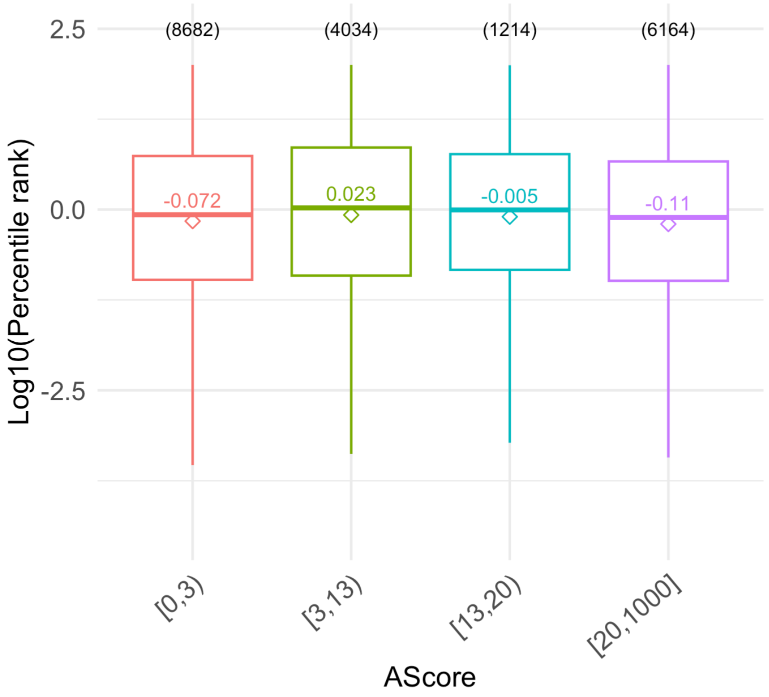


**Fig. 1**. **NetMHCIIphosPan log10 transformed predicted percentile rank scores for different AScore values.** The boxplots show the median of the plotted log10 percentile rank scores for each AScore interval. The center line inside the box also indicates the median, the mean is represented as an empty diamond and the box covers the interquartile range. The whiskers extend to, at most, 1.5-fold of the interquartile range. Above each boxplot, the number of ligands corresponding to each AScore interval is shown.

This analysis shows that for the different AScore intervals, the predicted percentile rank score distributions are highly overlapping. This result suggests that the NetMHCIIphosPan method is not sensitive to AScore values and does not benefit from a higher certainty in phosphosite localization.

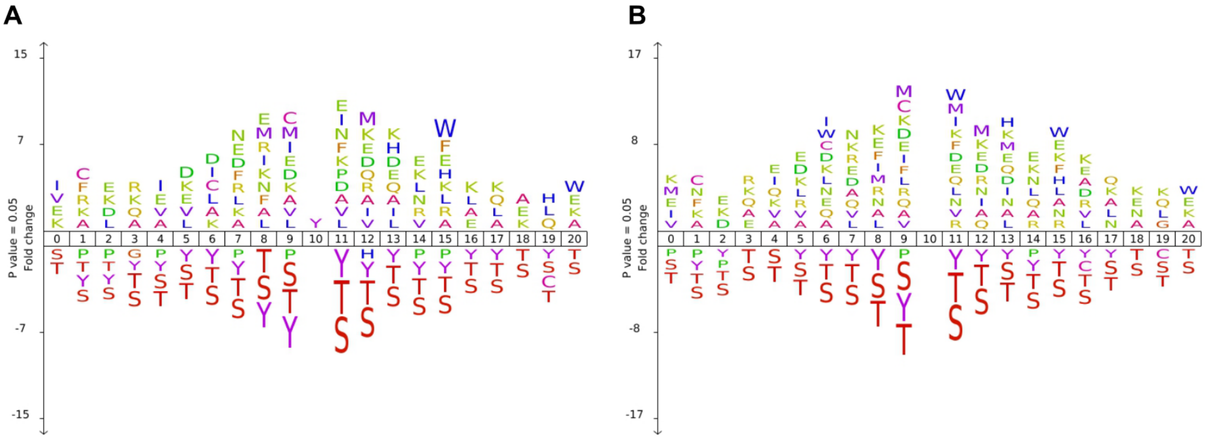

**Fig. 2. iceLogos depicting positional fold change in the frequency of amino acids between an "experimental" and a "reference" set of sequence-centered phosphosites with 10 flanking amino acids in the left and right regions** (the phosphosite is placed in position 10). Significantly enriched amino acids are represented above the x-axis, while depleted amino acids are shown below. The logos show a comparison between unambiguous (AScore > 13) and (**A**) ambiguous phosphosites (AScore <=13) or (**B**) S/T/Y potential acceptor sites, according to PEAKS. The number of sequences used to construct the logos: (**A**) N=5371 in the "experimental" set and N=10483 in the "reference" set; (**B**) N=5371 in the "experimental" set and N=23164 in the "reference" set.

Following, we constructed logos to visually inspect the difference in sequence composition between unambiguous (AScore > 13) and ambiguous phosphosites (AScore <=13), as annotated by PEAKS. **Fig. 2** consists of iceLogos[2] illustrating the positional fold change in the frequency of amino acids between these two sets of sequence-centered phosphosites (A), with 10 flanking amino acids in the left and right regions. Enriched amino acids are represented above the x-axis, while depleted amino acids are shown below. The logo in (A) (and similarly in (B), where the comparison is made with S/T/Y acceptor sites) shows a marked depletion of S/T/Y amino acids in both the left and right phosphosite flanking regions. This pattern is expected, as the absence of nearby acceptor sites favors precise phosphosite annotation by increasing the number of site-determining ions that could potentially differentiate two neighboring phosphosites. This is captured, by definition, in the AScore metric. On the contrary, there is no clear pattern of enriched amino acids in the proximity of the identified phosphosite, which suggests, as observed by Solleder *et al.*[3], that phosphosites in HLA-II ligandome emerge from a more diverse repertoire of kinases compared, for instance, with the previously studied phosphosites in HLA-I ligandome[4,5] Therefore, these observations, in line with the results shown in **Fig. 1**, indicate that models would not benefit from being trained on the subset of high-confidence phosphosites.

All the same, we inspected this in detail by comparing the five-fold cross-validation performance of a model trained and tested on all phosphopeptides reported by PEAKS, on phosphopeptides with AScore>13 (cutoff as defined in [6]) and phosphopeptides with AScore<=13. Again, for phosphopeptides with more than one phosphosite, the median AScore was calculated. Around 36% of all phospholigands have AScores>13 and the remaining 64% have AScores<=13 (see **Fig. 3**). Models were trained with the NNAlign_MA framework[7], using a five-fold cross-validation scheme. Training and test data were prepared in the same manner as mentioned before for the models shown in the main results section "Enhanced Predictive Models for Phospholigand Binding" (for more details refer to Materials and Methods: "Training datasets", "Data partitioning" or "Model training"). After splitting positive instances from each partition according to high/low AScore values, negatives were randomly sampled to preserve their uniform peptide length distribution. This implies that training datasets are of different sizes for the three models shown in **Fig. 3**. Nonetheless, model performances are comparable as the two test datasets ("AScore<=13" and "AScore>13") are consistent across all model evaluations.

A first observation from this result is that the model trained on the high AScore peptides achieves higher performance on the high AScore test data compared to the low AScore subset and that this trend is reverted for the model training on the low AScore data. However, the results also indicate that the model trained with all phosphopeptides ("Phos-only") performs better (or on par) across all metrics compared to the models trained only on phosphopeptides with AScore>13 ("Phos-only (AScore>13)") or on phosphopeptides with AScore<=13 ("Phos-only (AScore<=13)"). This result thus aligns with the finding in **Fig. 1**, suggesting that the modeling does not benefit from being focused on data with high phosphosite localization accuracy according to PEAKS.

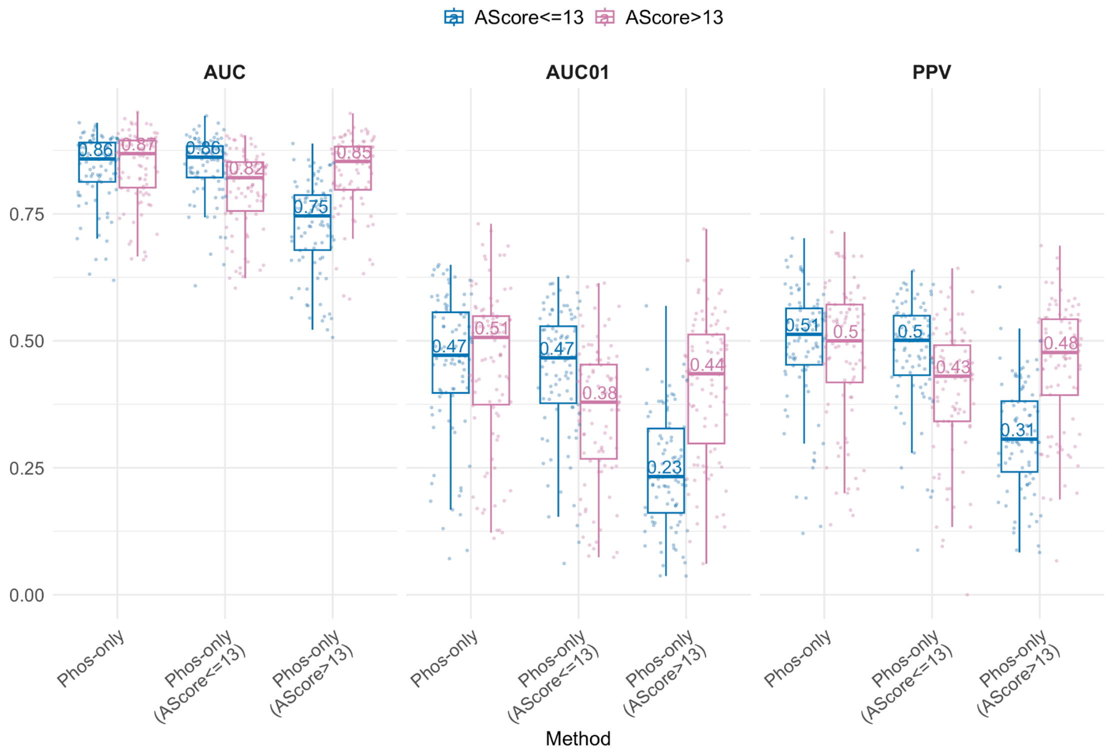


**Fig. 3. Five-fold cross-validation performance of the methods trained with phospholigands identified by PEAKS.** Performance metrics are reported for two subsets of the test data, phosphopeptides with AScore>13 and the remaining with AScore<=13. The "Phos-only" method was trained on all reported phosphopeptides, while "Phos-only (AScore>13)" was trained on phosphopeptides with AScore>13 and "Phos-only (AScore<=13)" was trained on phosphopeptides with AScore<=13. Performance metrics included are AUC, AUC0.1 (AUC integrated up to a false positive rate of 10%), and PPV (Positive Predictive Value, calculated as described in Materials and Methods). Performance is evaluated per dataset from the concatenated test sets (for details see Materials and Methods). The center line inside the box indicates the median of the plotted metric,

with its corresponding numerical value displayed above, and the box covers the interquartile range. The whiskers extend to, at most, 1.5-fold of the interquartile range. The individual data points in the jitter plots correspond to different cell lines.

Another strategy to assess the reliability of the identified phosphosites would be mapping phosphopeptides to phosphoproteomics databases. Here, we searched for all the phosphopeptides in our training database in PhosphoSitePlus[8]. This procedure recovered only 5.2% (881/16943) of the total unique phosphopeptide sequences in our training dataset. Furthermore, as the mentioned database compiles results from published papers and public MS datasets without applying any quality control on the reported phosphosites, we employed the compiled set of human phosphosites given by Kalyuzhnyy *et al.*[9]. Here, the authors curated all STY sites in the human proteome that have at least 5 instances of phosphorylation evidence in both PhosphoSitePlus and PeptideAtlas, creating a "gold standard" database of human phosphorylation sites. Only 4.3% (729/16943) of the total unique phosphopeptide sequences in our training dataset mapped to this "gold standard". Given these low recovery rates, we dismissed the strategy of filtering highly reliable phosphopeptides based on their presence in phosphoproteomics databases.

Next, we investigated if prediction tools could be applied to assess the reliability of the MS-identified phosphosites. Our first approach consisted of predicting phosphosites with NetPhosPan[10], a deep convolutional neural network (CNN) model that allows for the prediction of phosphosites in a "generic" or pan-receptor mode, leveraging information across a wide range of protein kinases. Alternatively, we also employed PhosphoLingo[11]. This recently developed tool employs protein language model embeddings instead of one-hot encodings to represent peptide sequences and feed them into a CNN model. Most interestingly, the PhosphoLingo study revealed that current phosphosite prediction methods inadvertently capture patterns related to the digestion specificity of MS protease enzymes. As a result, models tend to overestimate the accuracy of predictions for *de novo* phosphosites, influenced by these technical artifacts.

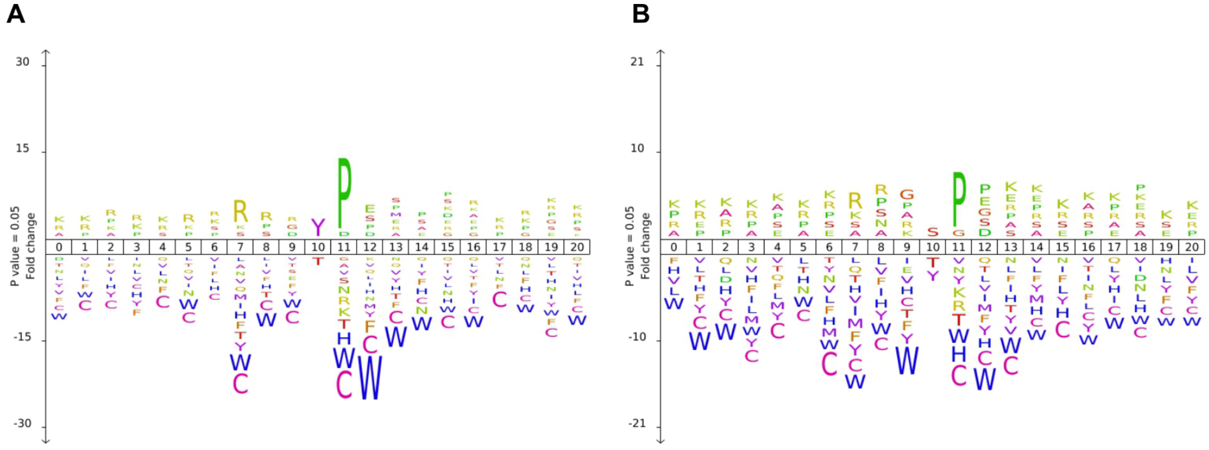


**Fig. 4. iceLogos depicting positional fold change in the frequency of amino acids between an "experimental" and a "reference" set of sequence-centered phosphosites with 10 flanking amino acids in the left and right regions** (the phosphosite is placed in position 10). Significantly enriched amino acids are represented above the x-axis, while depleted amino acids are shown below. The logos show a comparison between high vs. low-scoring phosphosites according to (**A**)

NetPhosPan and (**B**) PhosphoLingo predictions (cutoff was set at a score of 0.2). Number of sequences used to construct the logos: (**A**) N=2906 positives and N=11771 negatives; (**B**) N=3965 positives and N=13626 negatives.

Considering all these caveats, we predicted our training dataset phosphopeptides with the two described methods. Setting the (somewhat arbitrary) cutoff score at 0.2, 19.8% of all identified phosphosites were considered positive according to NetPhosPan, and 29% according to PhosphoLingo. **Fig. 4** consists of iceLogos[2] illustrating the positional fold change in the frequency of amino acids between two sets of sequence-centered phosphosites, with 10 flanking amino acids in the left and right regions. Enriched amino acids are represented above the x-axis, while depleted amino acids are shown below. In this case, we compared high vs. low-scoring phosphosites according to NetPhosPan (A) and PhosphoLingo (B) (cutoff was set at a score of 0.2). Logos (A) and (B) appear very similar, with a marked enrichment in P at position 11 (+1), denoting the preponderance of proline-dependent kinase phosphorylation patterns amongst high-scoring peptides. This motif typically corresponds to the family of cyclin-dependent kinases (CDKs), amongst others. We can also observe an enrichment of R (and also K in a smaller proportion) at positions 7 and 8 (-3 and -2 respectively). This motif is associated with calmodulin-dependent protein kinases (CAMKs), along with others[10,12] Inspecting the logos in **Fig. 4**, we observe that the enrichment signals in high vs. low-scoring predicted phosphosites (A and B) hold no relation to what is shown for unambiguous vs. ambiguous phosphosites reported by PEAKS (**Fig. 2**). We hypothesize that phosphosite predictors are biased towards cytosolic/nuclear protein kinase phosphorylation motifs since most of the proteins identified in phosphoproteomics experiments, and therefore employed to train these methods, derive from these subcellular compartments. Indeed, we suspect this is the reason why very few of the total identified phosphosites were found in public databases.

To further investigate this, we predicted the subcellular localization for the source proteins of our phosphopeptide training dataset with DeepLoc-2.0[13], classifying proteins in three subsets according to their localization: cytosolic (nucleus, cytoplasm, mitochondria and peroxisome), endolysosomal (extracellular, endoplasmatic reticulum, Golgi apparatus, lysosome/vacuole and cell membrane) and ambiguous. **Fig. 5** demonstrates that the higher the predicted PhosphoLingo score, the greater the fraction of phosphopeptides derived from cytosolic proteins. This implies that the predictor is heavily biased towards identifying cytosolic protein kinase motifs. Considering that our predictions indicate that more than 60% of phospholigands are derived from proteins located in the endolysosomal pathway, which is expected for natural HLA class II ligands, this result suggests that phosphosite predictors are not reliable tools to assess precise phosphosite localization.

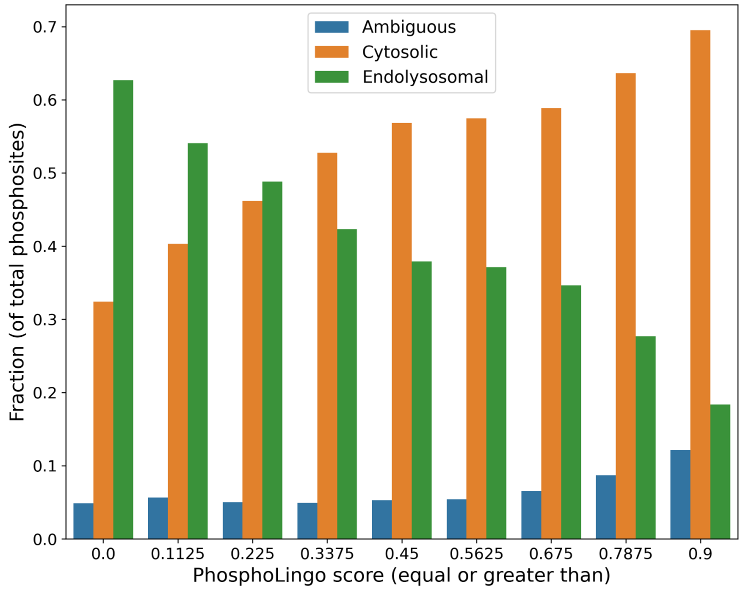


**Fig. 5**. **Subcellular localization of phospholigand source proteins was evaluated as a function of their predicted phosphosite scores.** The bars on the plot correspond to different thresholds for phosphosite scores. For example, the first set of bars (on the far left of the x-axis) includes all source proteins without any filtering (phosphosite scores ≥ 0). The second set of bars represents source proteins with phosphosites having predicted scores ≥ 0.1125, and so on. DeepLoc-2.0 was employed to predict protein subcellular localization, classifying proteins into three subsets: cytosolic (nucleus, cytoplasm, mitochondria, and peroxisome), endolysosomal (extracellular, endoplasmatic reticulum, Golgi apparatus, lysosome/vacuole and cell membrane) and ambiguous (more than one predicted compartment). PhosphoLingo was used to predict phosphosites.

(12) Johnson, J. L.; Yaron, T. M.; Huntsman, E. M.; Kerelsky, A.; Song, J.; Regev, A.; Lin, T.-Y.; Liberatore, K.; Cizin, D. M.; Cohen, B. M.; Vasan, N.; Ma, Y.; Krismer, K.; Robles, J. T.; van de Kooij, B.; van Vlimmeren, A. E.; Andrée-Busch, N.; Käufer, N. F.; Dorovkov, M. V; Ryazanov, A. G.; Takagi, Y.; Kastenhuber, E. R.; Goncalves, M. D.; Hopkins, B. D.; Elemento, O.; Taatjes, D. J.; Maucuer, A.; Yamashita, A.; Degterev, A.; Uduman, M.;

Lu, J.; Landry, S. D.; Zhang, B.; Cossentino, I.; Linding, R.; Blenis, J.; Hornbeck, P. V; Turk, B. E.; Yaffe, M. B.; Cantley, L. C. An Atlas of Substrate Specificities for the Human Serine/Threonine Kinome. *Nature* 2023, *613* (7945), 759–766. https://doi.org/10.1038/s41586-022-05575-3.

(13) Thumuluri, V.; Almagro Armenteros, J. J.; Johansen, A. R.; Nielsen, H.; Winther, O. DeepLoc 2.0: Multi-Label Subcellular Localization Prediction Using Protein Language Models. *Nucleic Acids Res* 2022, *50* (W1), W228–W234. https://doi.org/10.1093/nar/gkac278.
